# A SARS-CoV-2 entry inhibitor trimerizes to lock the spike protein in a closed conformation

**DOI:** 10.1101/2025.11.19.689107

**Authors:** Huihui Mou, Bo Gao, Gang Ye, Lizhou Zhang, Divyasha Saxena, Fan Bu, Jyoti Vishwakarma, Shuo Zhou, Claudia Ruiz Bayona, Li Lin, Yulia Getmanenko, Janarjan Bhandari, Charles C. Bailey, Gogce C. Crynen, Claire E. Kitzmiller, Hao Li, Yuka Otsuka, Chao Wang, Lalit Batra, Stuart Weston, Matthew B. Frieman, David K. Meyerholz, Louis Scampavia, Timothy P. Spicer, Reuben S. Harris, Michael D. Cameron, Thomas D. Bannister, Jian Zheng, Michael Farzan, Fang Li, Hyeryun Choe

## Abstract

The SARS-CoV-2 spike protein binds its receptor ACE2 to initiate target-cell infection. To engage ACE2, at least one of the three receptor-binding domains (RBDs) of the spike must adopt the up orientation. Here we describe S22, a potent, bioavailable, and non-toxic inhibitor of BA.2 and all subsequent Omicron variants. Cryo-EM analyses showed that S22 assembled as a trimer in a previously uncharacterized pocket of the spike apex, stabilizing all three RBDs in the down orientation, thereby preventing ACE2 association. Binding studies, especially those using mixed S22-sensitive and -resistant spikes, imply a cooperative assembly of three S22 molecules with three RBDs, resulting in an unusually slow S22 off-rate. Consistent with its slow dissociation and favorable pharmacokinetics, S22 suppressed viral replication 100-fold in the lungs of XBB.1.5-infected mice. Thus, S22 potently inhibits Omicron entry through a distinct mechanism whereby a small compound assembles cooperatively as a trimer to stabilize spike in an inactive conformation.

## Introduction

The entry glycoprotein spike of severe acute respiratory syndrome coronavirus 2 (SARS-CoV-2, hereafter SARS2) mediates entry into host cells^1–3^. The spike is synthesized as a ∼180 kDa protein that assembles into noncovalent trimers in the endoplasmic reticulum. After trafficking to the Golgi apparatus, it is cleaved into the S1 and S2 subunits by the cellular protease furin^4,5^. S1 binds to the receptor angiotensin-converting enzyme 2 (ACE2). ACE2 engagement and subsequent proteolytic activation steps^6–8^ initiate a series of conformational changes in both subunits that ultimately lead to fusion of the viral and cellular membranes, introducing viral genetic material into the cytoplasm. The S1 subunit comprises four domains: N-terminal domain (NTD), receptor-binding domain (RBD), and C-terminal domains (CTD) 1 and 2^9–11^. In the prefusion conformation of the spike trimer, S1 wraps around S2, with the three RBDs forming the apex.

The SARS2 RBD, like that of the other human sarbecovirus SARS-CoV (SARS1), is an independently folding domain that binds ACE2 with high affinity^12,13^. The SARS1 RBD was first described and structurally characterized shortly after the emergence of SARS1 in the winter of 2002-2003^14,15^. Subsequent cryo-electron microscopy (cryo-EM) studies of the spike proteins from SARS1, the merbecovirus MERS-CoV (MERS), and SARS2 revealed two general orientations of the RBD^9,16,17^. When all three RBDs of the spike trimer are symmetrically arranged in the down position, ACE2-binding residues of the RBD are occluded, and the spike in this closed conformation is unable to bind the receptor or mediate viral entry. The spike can also assume an asymmetrical conformation in which one or more RBDs are positioned away from the spike protein body (e.g., in the up position), thereby exposing ACE2-binding residues. Spike trimers in this open state can bind ACE2. The RBD is also the primary target of neutralizing antibodies, some of which sterically hinder its conformational transition to an open conformation^18,19^. However, the neutralizing activity of many of these antibodies has been reduced or abrogated with the emergence of Omicron variants, whose spike proteins vary extensively from the ancestral SARS2. In addition to antibodies, several SARS2 entry inhibitors that target the spike protein have been described. These inhibitors can block receptor binding^20,21^ or interfere with the downstream conformational changes required for membrane fusion^22^. Again, the activity of many of these compounds diminished with the emergence of Omicron. None of these compounds has been shown to inhibit spike’s transition to the active conformation or to revert the active conformation to the inactive one.

Here we identify a small molecule, S22, that binds and trimerizes within the apex cavity of the SARS2 spike trimer. S22 strongly inhibits viral entry in both pseudotyped virus and live virus assays, assembles as a trimer in a previously uncharacterized spike protein pocket, locks the RBDs in their down position as confirmed by conformation-specific antibodies and cryo-EM studies, and exhibits unusually slow dissociation as measured by surface plasmon resonance (SPR). In infected animals, S22 treatment reduces peak lung viral loads by two orders of magnitude and diminishes virus-induced pulmonary inflammation. These findings describe a self-interacting compound with an ability to trimerize, identify a druggable site in the coronaviral spike protein, and show that an assembled compound trimer can lock spike trimers in an inactive state to potently inhibit viral entry.

## Results

### High-throughput screening identified S22 as a SARS-CoV-2-specific entry inhibitor

We conducted three parallel high-throughput screens for viral glyprotein-specific entry inhibitors, in which two different pseudotyped viruses (PVs), each encoding either firefly luciferase (FLuc) or NanoLuc luciferase (NLuc), simultaneously infected cells (**Fig. 1a**)^23^, Simultaneous screening for selective SARS-CoV-2, Lassa, and Machupo virus entry inhibitors. This dual-reporter design enabled simultaneous monitoring of two viral entry events in a single well, allowing each virus to serve as a control for the other. This approach minimized experimental artifacts and eliminated inhibitors that act on cellular pathways. Using combinations of PVs bearing the entry proteins of Lassa virus (LASV), Machupo virus (MACV), and SARS-CoV-2 (SARS2, Omicron variant BA.5), we comprehensively screened a library of ∼665,000 drug-like small molecules^24^ and identified 50 compounds that specifically inhibited SARS2 entry with IC_50_ values below 10 μM and minimal or no cytotoxicity. We prioritized 21 candidates and further validated them (**Extended Data Table 1**), using the LASV entry inhibitor L2, the MACV entry inhibitor M8, and the coronavirus entry inhibitor hydroxychloroquine (HCQ) as controls (**Extended Data Fig. 1**). Among these, two compounds, S22 and S23, were selected for further investigation. S22 neutralized only SARS2, while S23 neutralized both SARS-CoV (SARS1) and SARS2. Neither compound inhibited PVs expressing arenaviral glycoprotein complexes (GPCs) of LASV and MACV.

**Fig. 1.**
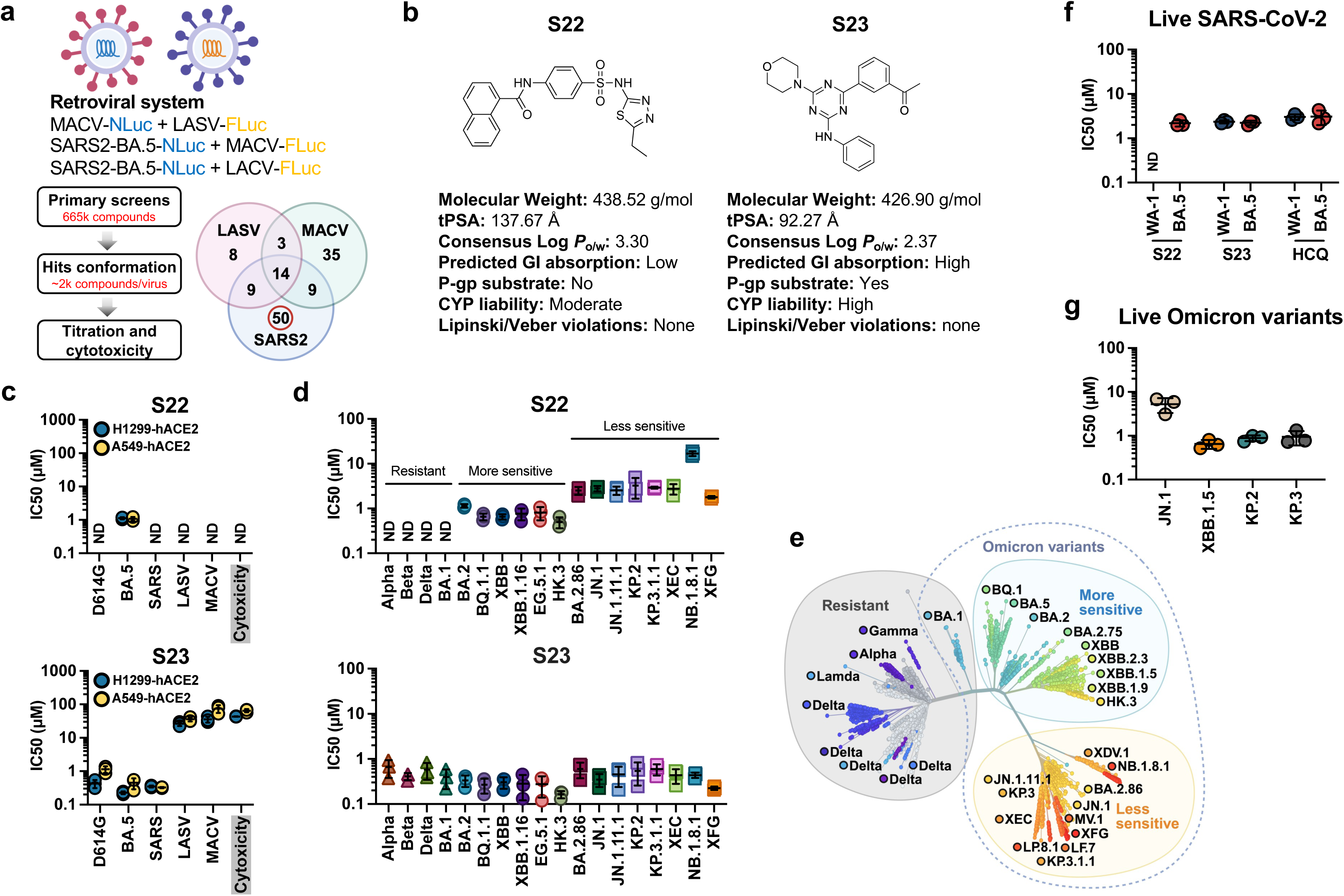
S22 is a SARS-CoV-2-specific entry inhibitor that neutralizes BA.2 and subsequent Omicron variants. **a**, Schematic of the paired entry assays and high-throughput screening (HTS) workflow used to identify entry inhibitors of Lassa virus (LASV, Josiah strain), Machupo virus (MACV, Carvallo strain), and SARS-CoV-2 (SARS2, BA.5 variant). Each virus pair includes two pseudotyped viruses (PVs) carrying two different reporters, firefly luciferase (FLuc) or NanoLuc luciferasae (NLuc), to enable simultaneous monitoring of dual infections in a single well. The Venn diagram summarizes the hit overlap. **b**, Structures of S22 and S23, and their representative drug-like properties predicted by SwissADME. **c**, Half-maximal inhibitory concentration (IC_50_) values of S22 and S23 against indicated PVs and cytotoxicity were determined in H1299-hACE2 cells (blue dots with black outlines) or A549-hACE2 cells (yellow dots with black outlines). Curves are shown in **Extended Data Fig. 2b**. **d**, S22 and S23 were assessed against the PVs bearing the indicated SARS2 spikes in H1299-hACE2 cells. Neutralization curves are shown in **Extended Data Fig. 2c**. **e**, SARS2 evolution correlates with the neutralizing activity of S22 against SARS2 variants shown in **d**. An unrooted phylogenetic tree was generated using Nextstrain/ncov based on 3334 genomes sampled globally from December 2019 to August 2025, with branch lengths indicating genetic divergence. The resulting tree was modified using BioRender. SARS2 variants shaded light gray are resistant to S22, light blue shaded variants are neutralized by S22 with higher potency, and yellow shaded variants are neutralized with comparatively lower potency. **f**, Antiviral activities of S22, S23, and hydroxychloroquine (HCQ) against authentic SARS2 ancestral WA-1 and BA.5 strains were assessed in Vero E6-hACE2 cells. Neutralization curves are shown in **Extended Data Fig. 2d**. **g**, S22 was assessed for its activity against additional authentic SARS2 variants. Neutralization curves and IC_50_ values are shown in **Extended Data Fig. 2e**. Data in **c** and **d** are representative of three independent experiments (n = 2 per concentration), and **f** and **g** include four replicates per concentration. Data are presented as mean ± s.d; ND, no detectable inhibition.

S22 and S23 are heteroaromatic small molecules predicted to have favorable drug-like properties (**Fig. 1b**). S23 is structurally similar to apilimod, a PIKfyve kinase inhibitor reported to block the entry of various viruses, including Ebola virus (EBOV) and SARS2, by perturbing endosomal trafficking of internalized viral particles^25^. Because apilimod, and by structural analogy likely S23, target a cellular pathway, we focused on S22, which was noncytotoxic, and employed S23 as a control. Antiviral profiling revealed the distinct breadth of these compounds: S23 inhibited SARS2-D614G, BA.5, and SARS1, and MERS-CoV (MERS) PVs, but not the arenavirus GPC controls, whereas S22 only neutralized the Omicron variant BA.5 (**Fig. 1c** and **Extended Data Fig. 2a,b**). Evaluation of additional SARS2 variants showed that S22 selectively inhibited BA.2 and later Omicron subvariants, including XFG, the most prevalent circulating strain (**Fig. 1d** and **Extended Data Fig. 2c**). In addition, we observed that the neutralizing activity of S22 against SARS2 variants is cluster-dependent. S22 was more potent against BA.5-related variants (“More sensitive” cluster in **Fig. 1e**, with average IC_50_ values ranging from 0.49-1.1 μM) than against JN.1-related variants (“Less sensitive” cluster, with average IC_50_ values ranging from 1.8-16.8 μM). This lineage-dependent pattern was not observed with S23.

S22 and S23 were further tested against authentic SARS2 isolates, with HCQ included as a control. Consistent with PV assays, S23 and HCQ neutralized the ancestral WA-1 and BA.5 variants, whereas S22 specifically inhibited Omicron variants emerging after BA.1 (**Fig. 1f,g** and **Extended Data Fig. 2d,e**). The IC_50_ values of S22 were 2.2 μM for BA.5, 0.66 μM for XBB.1.5, 3.8 μM for JN.1, 0.82 μM for KP.2, 0.90 μM for KP.3, respectively. These results show that S22 is a SARS2-specific inhibitor that neutralizes BA.2 and subsequent Omicron variants.

### S22 blocks RBD-ACE2 interaction

To define the stage of viral entry targeted by S22, we performed time-of-addition experiments (**Extended Data Fig. 3a**). Unlike the control compound S23, which remained active throughout the entire entry process, S22 was effective only when added before the unbound virus was removed in a wash step, indicating that it blocked viral attachment. Consistent with its antiviral profile, S22 induced a dose-dependent increase in melting temperature (T_m_) of the XBB.1.5 and BA.5 spikes in the nano differential scanning fluorimetry (nano DSF) assays but did not increase the T_m_ of the control D614G spike (**Extended Data Fig. 3b**). Collectively, these data show that S22 associates directly with the spike to prevent attachment of the virus to the cell.

To determine if S22 inhibited spike engagement with the receptor ACE2, we expressed immunoadhesin forms of spike receptor-binding domains (RBD-Fcs) of two S22-sensitive variants, JN.1 and XBB, as well as those of two resistant viruses (WA-1 and MERS-CoV). We then assessed their binding to cell-surface ACE2 in the presence of S22, using the LASV entry inhibitor L2 as a control (**Fig. 2a** and **Extended Data Fig. 3c,d**). Consistent with its neutralization profile, S22, but not L2, inhibited JN.1- and XBB-RBD binding to ACE2 in a dose-dependent manner, while showing no effect on WA-1 (ancestral) RBD binding to ACE2 or MERS RBD binding to its receptor dipeptidyl peptidase 4 (DPP-4)^26^.

**Fig. 2.**
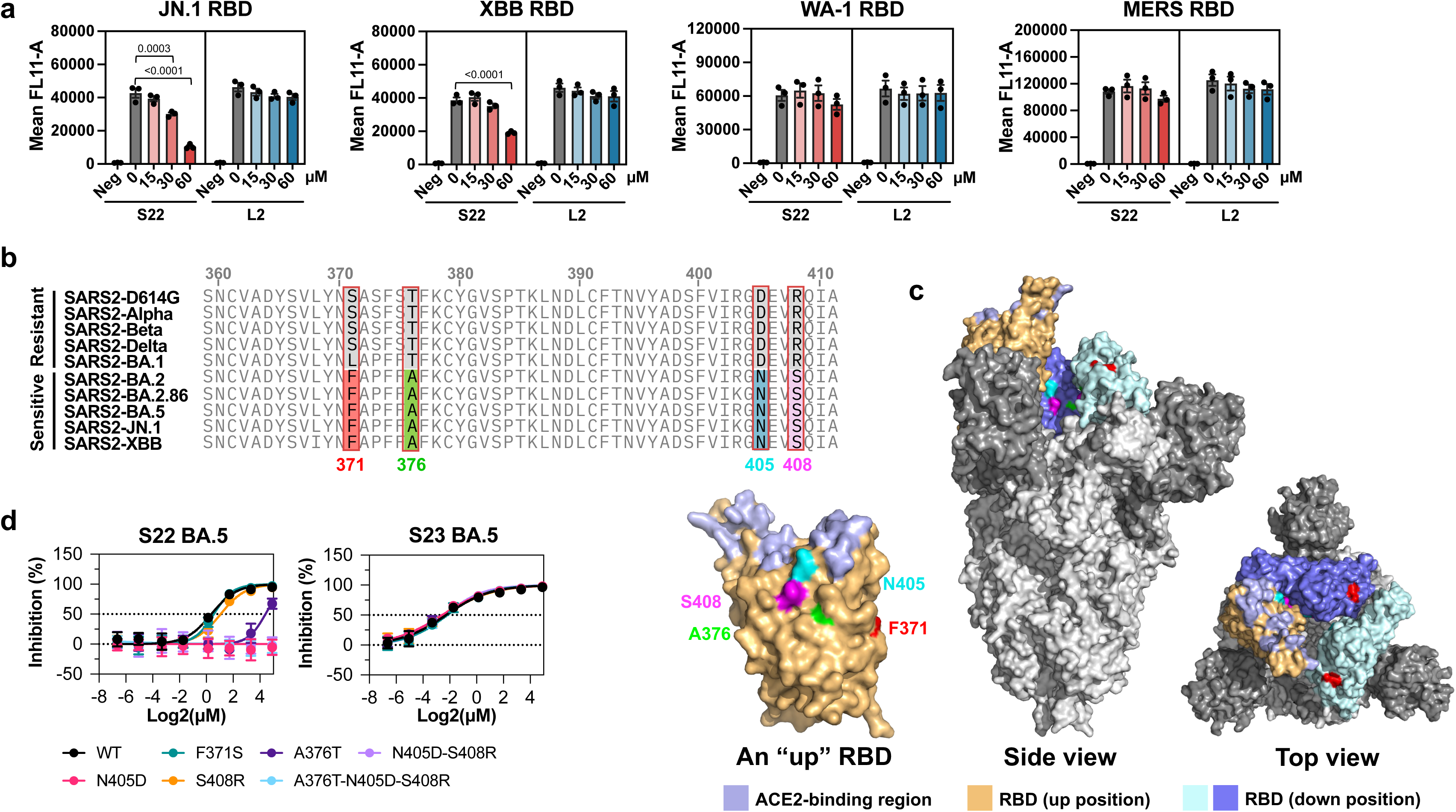
S22 blocks Omicron RBD-ACE2 interactions. **a**, The immunoglobulin form of the spike receptor-binding domains (RBD-Fc) from SARS2 Omicron variants (JN.1 and XBB), the ancestral WA-1 strain, and MERS-CoV were pre-incubated with the indicated concentrations of S22 or a control compound L2 at room temperature for 30 minutes before being added to H1299-hACE2-hDPP4 cells, and detected using APC-conjugated anti-human Fc antibody. Data represent mean fluorescence intensity (MFI) from three independent experiments, with error bars indicating s.d. Representative histograms are shown in **Extended Data Fig. 3d**. Background-corrected values were analyzed with generalized linear model, and post-hoc Dunnett’s test was used to assess statistical significance. Adjusted *p*-values are reported only for comparisons reaching statistical significance. **b**, Sequence alignment of spike RBDs from S22-sensitive (BA.2, BA.2.86, BA.5, JN.1, and XBB) and S22-resistant (D614G, Alpha, Beta, Delta, and BA.1) variants identified four residues (boxed in red) that differ between the two groups. **c**, Mapping of the four RBD residues identified in **b** onto the BA.5 spike (PDB ID: 8GTO). The middle and right panels show the side and top views of the spike trimer with one RBD in the up conformation. The left panel shows the inward-facing surface of the BA.5 RBD in the up conformation, with the four residues and ACE2-binding region indicated. **d**, S22 activity was assessed against PVs bearing the wild-type (WT) BA.5 spike or the mutants with the indicated mutations from S22-resistant spikes, either alone or in combination. S23 was included as a control. Data represent mean values from three independent experiments (n = 2 per concentration), with error bars indicating s.d.

Sequence alignment of the RBDs from five S22-sensitive and five resistant variants identified four residues of interest: A376, N405, and S408 located on the inner face of the RBD, positioned near the trimer axis of the closed spike, and F371 positioned at the outer face (**Fig. 2b,c**). Introducing residues from resistant variants into the BA.5 spike individually or in combination showed that N405D and, to a lesser extent, A376T impaired spike’s sensitivity to S22, but did not affect its sensitivity to S23 (**Fig. 2d**). These substitutions consistently conferred similar resistance when introduced into the spike of additional S22-sensitive variants (BA.2, JN.1, and XBB, **Extended Data Fig. 3e**). Together, these findings show that S22 directly blocks RBD-ACE2 association, which can be attenuated by changes in the inner face of the RBD.

### S22 assembles into a trimer that locks the spike in a closed conformation

To define the binding mode of S22, we determined the cryo-electron microscopy (cryo-EM) structure of the BA.5 spike in complex with S22. Size-exclusion chromatography (SEC)-purified BA.5 spike ectodomain in the prefusion conformation (0.8 mg/mL) was incubated with or without 100 μM S22 and vitrified on copper grids for data collection. Analysis of the resulting structures revealed that co-incubation with S22 altered the ratio of spike proteins in the open conformation (one RBD up) to those in its closed conformation (all RBDs down). In the absence of S22, 47% of the BA.5 spikes were observed in the open conformation (3.15 Å resolution), while 53% were found in the closed conformation (2.83 Å resolution, **Fig. 3a** and **Extended Data Fig. 4**). In contrast, when bound by S22, all BA.5 spikes were observed in the closed conformation (2.66 Å resolution). Notably, compared to the closed apo structure, the S22-BA.5 spike complex displayed an additional density within the central cavity of the spike trimer, which is well fitted by three S22 molecules assembled into a trimeric form (**Fig. 3b**). The S22 trimer was stabilized by strong hydrophobic contacts and hydrogen bonds formed between neighboring S22 molecules (**Fig. 3c**), simultaneously engaging the RBDs of all three protomers to lock the spike in its closed state (**Fig. 3d**).

**Fig. 3.**
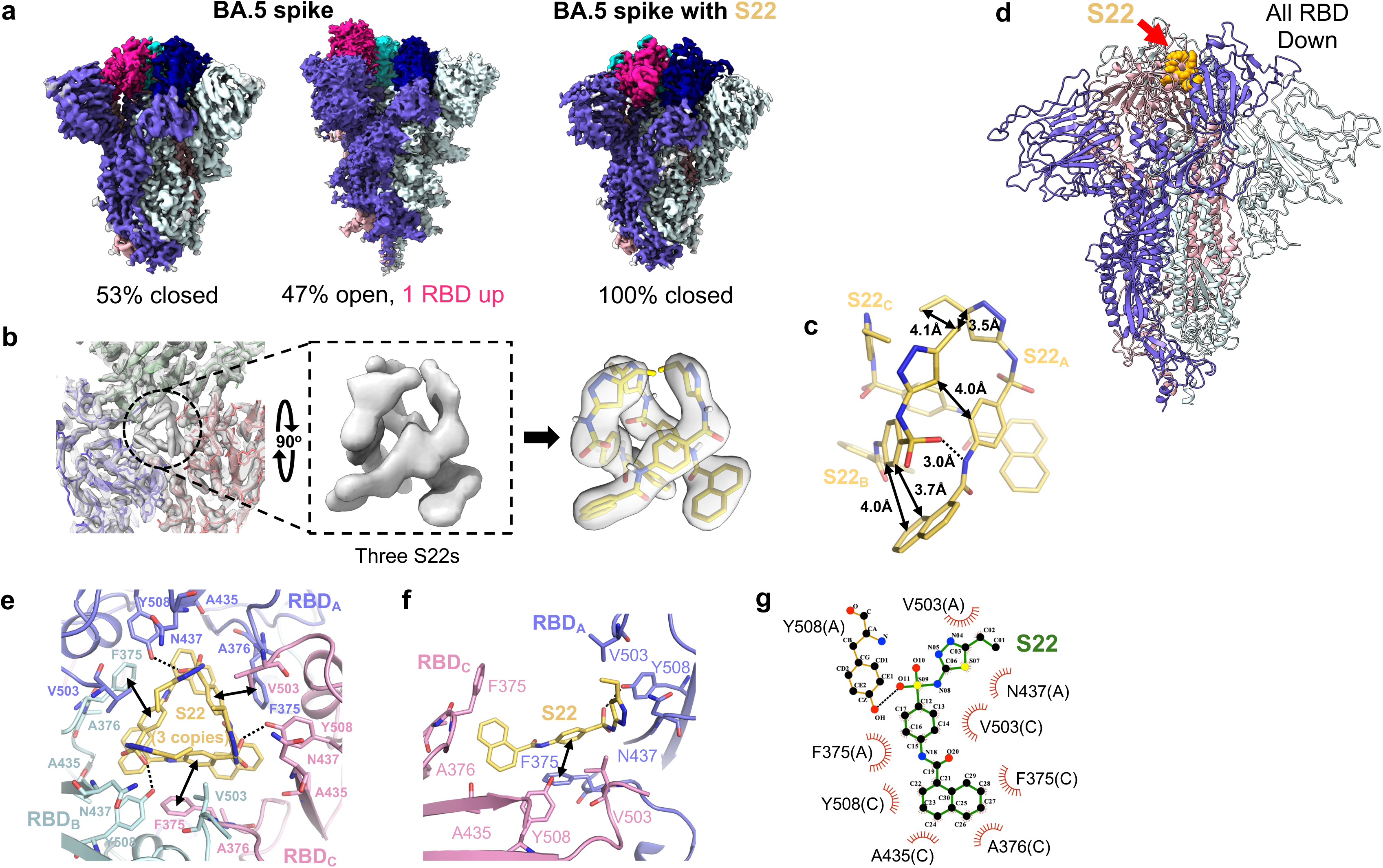
S22 assembles into a trimeric complex in the central cavity of the spike trimer, locking the spike in a closed conformation. **a**, Cryo-EM reconstructions of the BA.5 spikes with and without bound S22. The BA.5 apo protein adopts two conformations: closed (all RBDs down, left) and open (one RBD up, middle), with the proportion of particles in each state indicated below. With S22 binding, all spike proteins adopt the closed conformation (right). Each protomer is colored using a single-color gradient, with the darker shade highlighting the corresponding RBDs. **b**, Top view of the BA.5 spike-S22 complex (left) reveals additional electron density within the central cavity of the spike trimer. A zoom-in view shows this extra density (middle), which was fitted by three S22 molecules (right). **c**, Stick representation of S22 trimer to show the detained interactions between neighboring S22 molecules. **d**, Cartoon representation of the right panel of **a**, with the S22 trimer shown in yellow. **e**, All interacting side chains of spike residues are shown as sticks and labeled. **f**,**g**, Detailed interactions between a single S22 molecule and two adjacent RBDs, shown in a cartoon representation generated by PyMOL (**f**) and shown in a schematic diagram generated by LigPlot^+^ (**g**). For **c, e**, and **f,** dashed lines denote hydrogen bonds, and double arrows indicate hydrophobic contacts.

Analysis of the interaction interface between the S22 trimer and the BA.5 spike identified six RBD residues (F375, A376, A435, N437, V503, and Y508) in direct contact with the S22 trimer (**Fig. 3e**). F375 from one spike protomer formed π-π interactions with S22, while Y508 from the same protomer forms a hydrogen bond. Each S22 monomer bridges two neighboring RBDs by interacting with four residues from one RBD and five from a second, thereby locking the three RBDs in their down orientations (**Fig. 3f,g**). Phylogenetic analysis of representative spike sequences showed that residues critical for S22’s activity are conserved across all neutralized variants (**Extended Data Fig. 5a**). F375 first appeared in BA.1, an S22-resistant Omicron lineage closely related to other S22-sensitive variants, and interacts with S22 through π-π stacking with its core benzene ring. Introducing the F375S mutation into the BA.5 spike abolished its sensitivity to S22 (**Extended Data Fig. 5b**). The residues T376 and D405 of the BA.1 spike attenuate or abolish BA.5’s sensitivity to S22 when introduced into its spike (**Fig. 2d**). Consistently, introducing T376A and D405N mutations conferred BA.1’s sensitivity to S22 (**Extended Data Fig. 5c**). Of note, although NB.1.8.1, like all Omicron variants tested other than BA.1, remained sensitive to S22, it was less sensitive than most other Omicron variants (**Fig. 1d**). This variant carries a unique A435S mutation that has not been observed in previous strains. Introducing A435S into the BA.5 spike reduced its sensitivity to S22 by approximately 4-fold, indicating its role in S22-mediated neutralization (**Extended Data Fig. 5d**).

We also experimentally probed S22-induced conformational changes by comparing the binding of recombinant proteins and SARS2 neutralizing antibodies to BA.5 spikes expressed on cell surfaces, with or without S22 pre-treatment (**Fig. 4a**). Four immunoadhesins were used: ACE2-Fc and N3113-Fc, which recognize only the RBD in the up conformation, SP1-77, which binds the RBD in both conformations, and the LASV GPC antibody 8.9F as a control (**Extended Data Fig. 6a**). Entry protein controls included the D614G spike and the LASV GPC. S22 treatment did not alter expression of any viral entry proteins, as indicated by comparable levels of FLAG-tag staining in treated and untreated cells (**Extended Data Fig. 6b**). Consistent with the structural studies, S22 binding prevented the RBD from transitioning to an up conformation, as indicated by 12- and 40-fold reductions of ACE2-Fc and N3113-Fc binding, respectively, to the BA.5 spike. In contrast, S22 did not affect SP1-77 binding, nor did it impact ACE2 or antibody binding to the D614G spike or LASV GPC. Collectively, these results support a mechanism by which S22 assembles as a trimer to lock the spike in its closed conformation. To our knowledge, this trimeric assembly and resulting entry-inhibition mechanism are unique to S22.

**Fig. 4.**
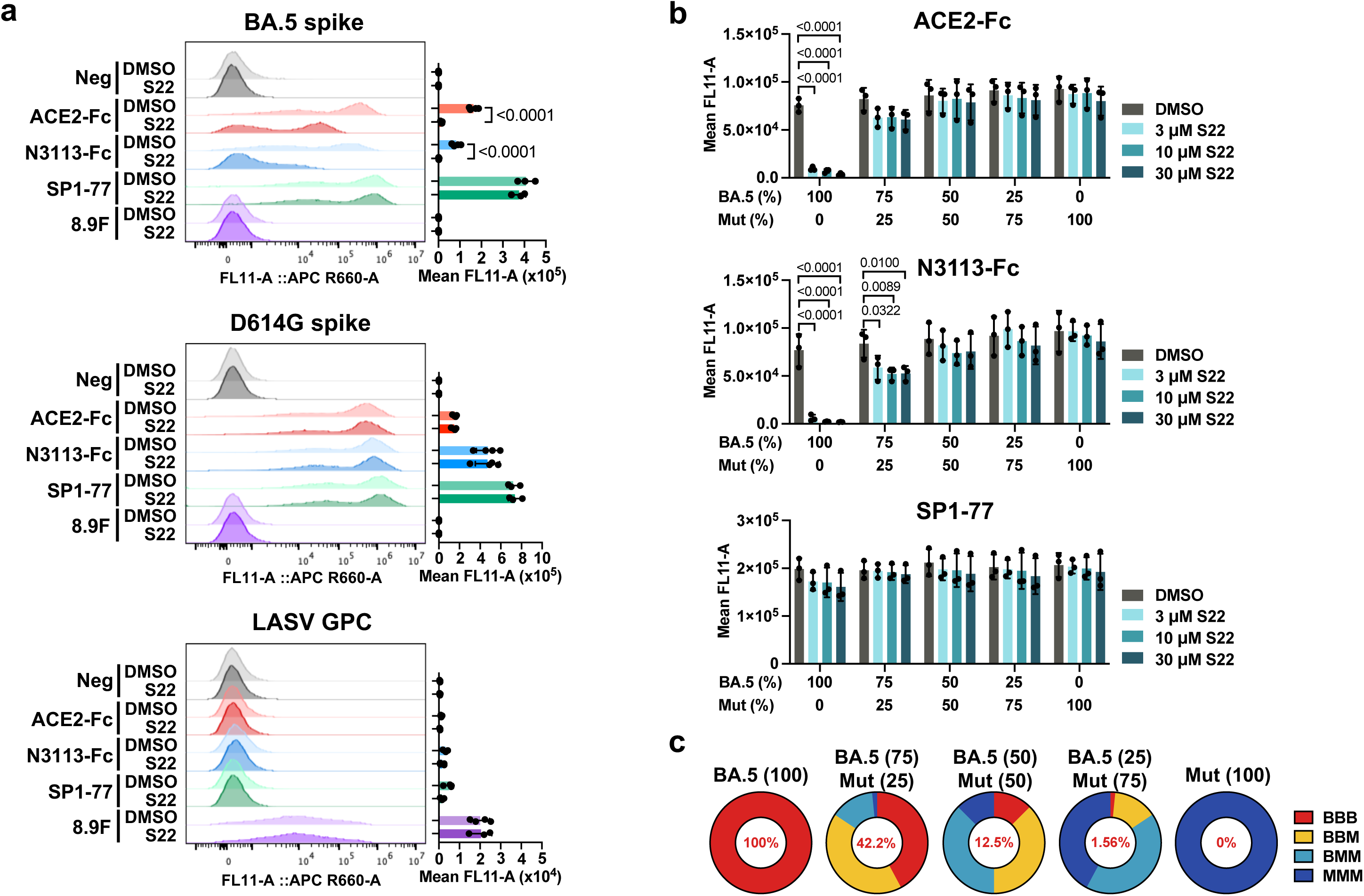
S22 cooperatively assembles into a trimer, locking the spike in the closed conformation. **a**, To assess S22-binding-induced conformational changes in the BA.5 spike, HEK293T cells expressing C-terminal FLAG-tagged entry proteins of BA.5, D614G, and LASV were treated with 30 µM S22 or DMSO for 1 h at 48 hour post-transfection (h.p.t.), fixed, and stained with ACE2-Fc or the indicated antibodies, followed by APC-conjugated anti-human Fc. Entry protein expression, treated with S22 or DMSO, was assessed by staining permeabilized cells with anti-FLAG antibody, as shown in **Extended Data Fig. 6b**. MFI values were square-root-transformed and analyzed using two-way ANOVA, followed by Sidak’s multiple comparisons test. **b**, To investigate whether S22 trimerization is required to induce conformational changes in the BA.5 spike, HEK293T cells were transfected with WT BA.5 spike (BA.5), the S22-resistant BA.5 F375S mutant (Mut), or mixtures of both at the indicated ratios. Cells were treated with S22 (3, 10, or 30 µM) or DMSO, then fixed and stained with ACE2-Fc or SARS2 neutralizing antibodies. MFI values were background-subtracted and analyzed using generalized linear model (gamma, loglink), and Dunnett’s test was used to assess statistical significance. **c**, The predicted proportion of homo and hetero spike trimers formed in each transfection condition in **b**, based on binomial probability, assuming random assembly of protomers. “B” and “M” in BBB, BBM, BMM, and MMM indicate WT BA.5 and F375 mutant spike protomer, respectively. For **a** and **b,** bars represent MFI from independent experiments (a: n = 4; b: n = 3), and error bars indicate s.d. Adjusted *p*-values are reported only for comparisons reaching statistical significance.

### Exceptionally slow dissociation of trimeric S22 from the spike

Structure analysis of the BA.5 spike-S22 complex suggests that cooperative assembly and trimerization are essential for S22’s activity. To investigate this possibility, we co-expressed WT BA.5 spike with its variant resistant to S22 (F375S; **Extended Data Fig. 5b**) at different ratios to generate heterotrimers. All combinations expressed comparably, as indicated by similar SP1-77 staining (**Fig. 4b**). Consistent with previous observations, S22 inhibited most of ACE2-Fc and N3113-Fc binding when all spikes were WT BA.5 homotrimers (**Fig. 4b,c**). In contrast, co-expression of 25% of the S22-resistant mutant spike, with an estimated 42.2% of WT homotrimers, restored approximately 80% of ACE2-Fc and N3113-Fc binding, indicating substantial attenuation of S22’s activity. S22 binding was nearly completely abrogated when 50% of co-expressed spikes were F375S, despite an estimated 12.5% of WT homotrimers. Similar results were observed in neutralization assays using PVs produced from mixed WT and F375S spikes (**Extended Data Fig. 6c**). Thus, even a single S22-resistant spike protomer of a trimer can impede S22’s binding and entry inhibition activity.

We next explored the binding kinetics of S22 to SARS2 spike proteins using surface plasmon resonance. Single-cycle kinetic analyses revealed biphasic dissociation of S22 from the XBB.1.5 and BA.5 spikes, one very fast and the other immeasurably slow, reflected by a progressive baseline increase with each cycle, consistent with retention of bound S22 (**Extended Data Fig. 7a,b**). The fast off-rate likely corresponds to S22 monomer dissociation, whereas the slow dissociation phase reflects stably bound S22 trimers. No baseline shift was observed for the D614G spike or EBOV GP. We further examined the dissociation kinetics of S22 from the BA.5 spike using a time-course flow cytometry assay (**Extended Data Fig. 7c**). Cells expressing BA.5 spikes were preincubated with or without S22 for one hour before medium replacement. When treated with 3 μM S22, gradual dissociation of S22 was evident from the time-dependent recovery of ACE2-Fc and N3113-Fc binding. In contrast, when treated with 30 μM S22, limited S22 dissociation was observed, consistent with stable engagement of S22 trimers with the BA.5 spike. This pattern was not observed for the D614G spike. We conclude that monomeric S22 rapidly disassociates from the spike, but once assembled, S22 trimers have an unusually slow off-rate. We further infer that this unusually slow off-rate results from the cooperative assembly of an S22 trimer with three RBDs.

### Single-dose S22 treatment inhibits SARS-CoV-2 replication in mice

Computational analyses using SwissADME predicted that S22, an amide derivative of the antibiotic sulfaethidole, possesses favorable drug-like properties (**Fig. 1b**). We also performed both in vitro and in vivo drug metabolism and pharmacokinetics (DMPK) studies. In these assays, S22 exhibited high in vitro metabolic stability with minimal CYP450 inhibition (**Extended Data Fig. 8a**) and displayed a plasma half-life of approximately three hours (**Extended Data Fig. 8b**). S22 reached a peak plasma concentration (C_max_) of 87 µM one hour after intraperitoneal administration and remained detectable 24 hours post-injection.

Given these favorable pharmacokinetic properties, we assessed the in vivo antiviral efficacy of S22 in an XBB.1.5-infected mouse model. C57BL/6 female mice (8- to 10-week-old) were intranasally inoculated with the virus (1 × 10^4^ PFU), and four hours post-infection, injected intraperitoneally with one dose of 40 mg/kg S22, 20 mg/kg of the SARS2 protease inhibitor ensitrelvir, or PBS control (**Fig. 5a**). Consistent with the previous observations^27,28^, XBB.1.5 infection caused no obvious clinical symptoms or weight loss over the 5-day study period (**Extended Data Fig. 8c,d**). S22 treatment nonetheless reduced lung viral titers by 100-fold at two days and approximately 20-fold at five days post infection. compared to PBS controls, as measured by TCID_50_ assays (**Fig. 5b**). Histopathological analyses further showed that both S22 and ensitrelvir mitigated pulmonary inflammation, indicated by no edematous lesions and lower pathology scores compared to PBS-treated mice (**Fig. 5c,d** and **Extended Data Fig. 8e,f**). To assess the potential for viral escape, we sequenced spike RNA from lung homogenates of mice two and five days post infection. (**Fig. 5e**). No consistent RBD mutations were observed, and no consistent mutations were detected in other parts of the spike in any group. Together, these findings demonstrate that S22 suppresses SARS2 replication in vivo and mitigates virus-induced lung pathology.

**Fig. 5.**
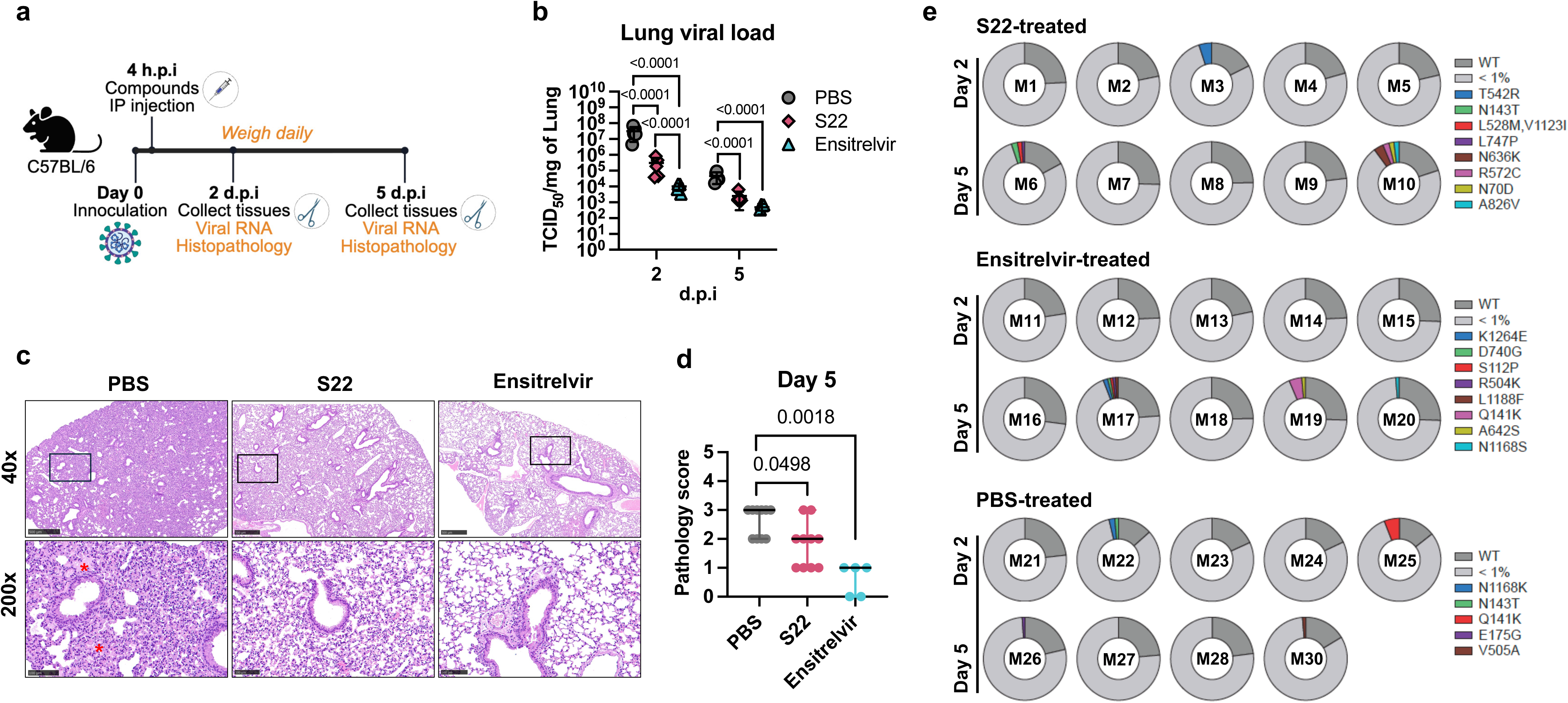
S22 inhibits viral replication and pulmonary inflammation in mice. **a**, Schematic of the therapeutic mouse study timeline. 8- to 10-week-old female C57BL/6 mice (n = 10 per group) were inoculated intranasally with 1 × 10^4^ PFU of SARS2 XBB.1.5. At 4 hours post infection (h.p.i.), mice were administered intraperitoneally with S22 (40 mg/kg), ensitrelvir (20 mg/kg), or PBS. Body weight was monitored daily as shown in **Extended Data Fig. 8c,d**. Lung tissues were collected on two and five days post infection (d.p.i.). for viral load quantification and histopathological analysis. Illustration was created with BioRender.com. **b**, Viral loads in lung tissues were measured by TCID_50_ assays. Data are presented as mean ± s.d. Lung viral load was log10-transformed and assessed for statistical significance by two-way mixed model ANOVA, followed by Šídák’s multiple comparisons test. **c,d**, Representative H&E-stained lung sections at both 40x and 200x magnifications (**c**) and corresponding pathology scores (**d**) at five d.p.i. Localized pulmonary edema, observed only in the PBS-treated group on day 5, is indicated by red stars. Day 2 images and pathology scores are shown in **Extended Data Fig. 8e,f**. Scale bars: 500 µm for 40× images; 100 µm for 200× images. Pathology scores were evaluated by ordinal logistic regression model, with significance determined by chi-square tests with Pearson’s correction. **e**, Spike sequences retrieved from lung homogenates were analyzed to monitor the emergence of escape mutations in vivo. Data are shown for all three treatment groups on two and five d.p.i. In each pie chart, the proportion of WT XBB.1.5 spike is shown in dark gray, low-frequency random variants (<1%) are grouped in light gray, and individual mutations with >1% frequency are indicated separately. For **b** and **d**, Adjusted *p*-values are reported only for comparisons reaching statistical significance.

## Discussion

We describe here the identification and characterization of the SARS-CoV-2 (SARS2) entry inhibitor S22 with several highly distinctive properties. It is potent against recently circulating Omicron variants. It assembles as a trimer, unique for an antiviral compound, at a previously uncharacterized and druggable spike cavity. Each S22 monomer in a trimer binds two RBDs, and each RBD engages two S22 molecules, stabilizing a tight, symmetrical spike locked in an inactive conformation. The cooperative nature of trimer assembly underlies the exceptionally slow off-rate of S22. This slow off-rate, favorable pharmacokinetic properties, and low cytotoxicity contribute to its potent in vivo anti-viral activity.

Other small molecules, such as linoleic acid, trans-retinoic acid, and S416, have been previously reported to stabilize the RBD-down spike conformation^29–32^. Each of these compounds binds independently, either to an individual RBD or at an RBD-RBD interface, within a hydrophobic pocket positioned away from the central cavity. In each case, their hydrophobicity precludes their potential for in vivo use. In contrast, S22 forms a trimer at the spike central cavity, and cooperative trimerization moves an RBD to a down state, as shown in cryo-EM. In the absence of S22, approximately half of soluble spike trimers adopt the open conformation with one RBD up. Upon S22 binding, all open spike trimers revert to their closed conformation, indicating S22 trimerization can drive the RBD from up to down orientation. The fully assembled complex of three S22s and three RBDs is highly stable, leading to undetectable dissociation three hours after binding to BA.5 spikes. We postulate that initial binding of the first S22 monomer is inefficient, but the affinity progressively increases as additional S22 molecules assemble at the binding pocket, because each subsequent S22 monomer makes contact with previous monomers as well as two RBDs. As the third S22 completes the trimer, it provides sufficient binding energy to move the up RBD to the down position, thereby locking the spike into the closed conformation. This unusual cooperative assembly process may also explain S22’s potent inhibition of viral infection in vivo, as it could remain associated with the spike in infected cells throughout the long window of spike assembly. Thus, S22 is distinguished from these previously reported compounds by its mechanism of inhibition, its unique active form as an assembled trimer, its potency, and its favorable pharmacokinetic properties.

S22 inhibits the entry of all characterized Omicron variants except BA.1, one of the first Omicron strains to emerge. The reason for the conservation of this cavity in post-BA.1 Omicron variants, despite continuous antigenic drift, is unclear. This region is under considerable immune pressure^33–38^ because the residues lining this cavity are exposed when the RBD is in the up conformation and accessible to neutralizing antibodies. The persistence of this cavity after several years of Omicron evolution suggests that its divergence from pre-Omicron strains confers some fitness advantage. The basis for this advantage remains unclear except that the residues in the cavity contribute to the balance between open and closed spike conformations, and they are proximal to the receptor-binding motif of the RBD, which directly engages ACE2. Our inability to observe a common or dominant viral escape pathway after five days of XBB.1.5 infection in mice with S22 treatment is consistent with the functional roles of the Omicron residues in this pocket.

The unique properties and mechanism of S22 may have other applications. Most immediately, compounds with a similar propensity to trimerize may be used to target other pathogenic coronaviruses. For example, the MERS-CoV spike undergoes similar conformational transitions from a closed, RBD-down state to an open, RBD-up state, and its RBDs form a cavity comparable to the one observed in the SARS2 spike^17,39^. More generally, compounds that cooperatively assemble at their binding sites could be designed to target multimeric proteins with symmetrical, hard to access pockets, including members of the TNFα superfamily, bacterial porins, and most viral entry proteins^40–45^. The trimeric core of S22 may serve as a scaffold for designing similar inhibitors with a propensity to trimerize.

In summary, we describe here an unusual mechanism of action for a small-molecule entry inhibitor whereby the viral entry protein is tightly locked in an inactive conformation incapable of binding its receptor. The compound effecting this mechanism, S22, does so by emulating the trimeric symmetry of its target protein spike to form multiple cooperative interactions that stabilize an assembled complex in its binding pocket, resulting in an unusually slow dissociation rate. S22 could thus serve as a model for a new class of self-assembling compounds that can target many cellular and microbial protein multimers.

**Extended Data Fig. 1.**
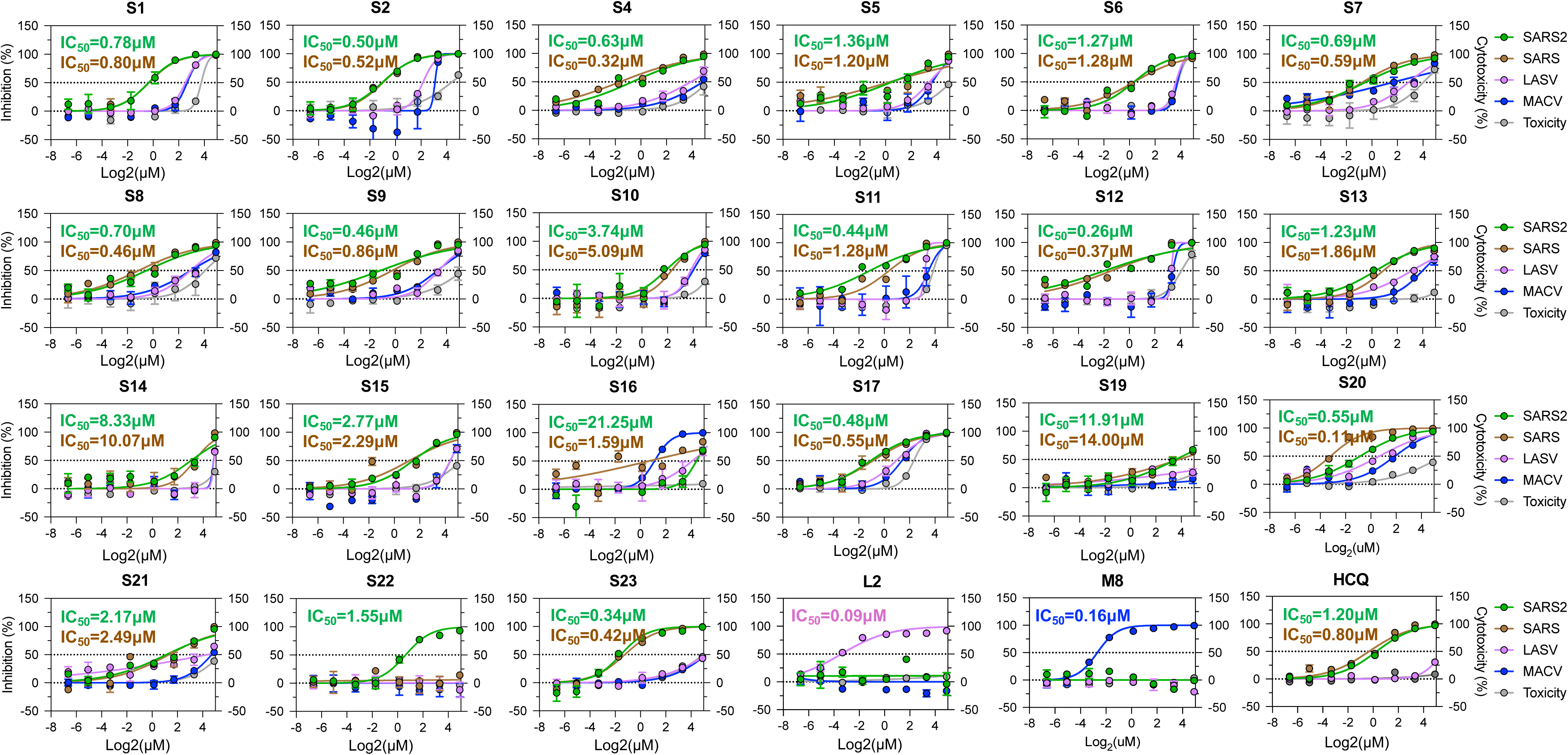
Validation of the SARS-CoV-2 entry inhibitors identified from HTS. 21 SARS2 entry inhibitors, prioritized from the 50 leads identified in the HTS, were evaluated for neutralizing activity against PVs bearing entry proteins of SARS2 (green), SARS-CoV (brown), LASV (pink), and MACV (blue) in H1299-hACE2 cells, using the LASV inhibitor L2, the MACV inhibitor M8, and HCQ as controls. Left Y-axis represents relative inhibition normalized to DMSO controls; Right Y-axis represents their cytotoxicity (grey). The data shown are representative of two independent experiments (n = 2 per concentration), with error bars indicating s.d. IC_50_s shown are color-coded to match their respective PVs.

**Extended Data Fig. 2.**
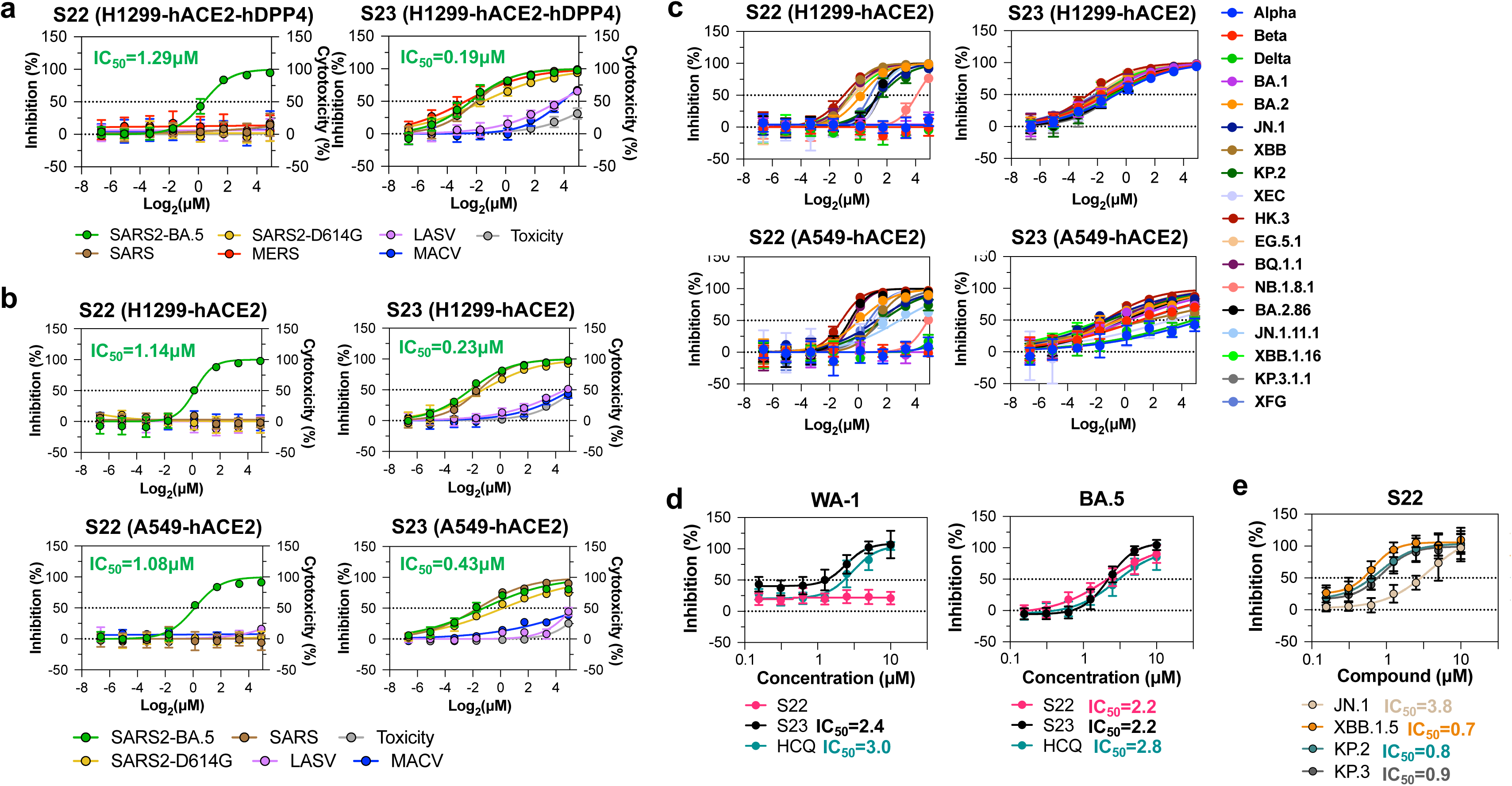
S22 neutralizes BA.2 and subsequent Omicron variants. **a**, S22 and S23 were assessed for their neutralizing activities against the indicated PVs and cytotoxicity in H1299-hACE2-hDPP4 cells. **b**, Neutralization curves of S22 and S23 against the indicated PVs in H1299-hACE2 and A549-hACE2 cells. The green values in **a** and **b** indicate S22 and S23’s IC_50_ against SARS2 BA.5. **c**, Neutralization curves of S22 and S23 against the indicated SARS2 PVs in H1299-hACE2 and A549-hACE2 cells. IC_50_ values from H1299-hACE2 cells are present in Fig. 1d. **d**, Neutralization curves of S22, S23, and HCQ against the authentic SARS2 ancestral WA-1 and BA.5 variants in Vero E6-hACE2 cells. IC_50_ values are present in Fig. 1f. **e**, Neutralization curves of S22 against the indicated authentic SARS-CoV-2 Omicron variants in Vero E6-hACE2 cells. IC_50_ values are present in Fig. 1g. Data are presented as mean ± s.d.

**Extended Data Fig. 3.**
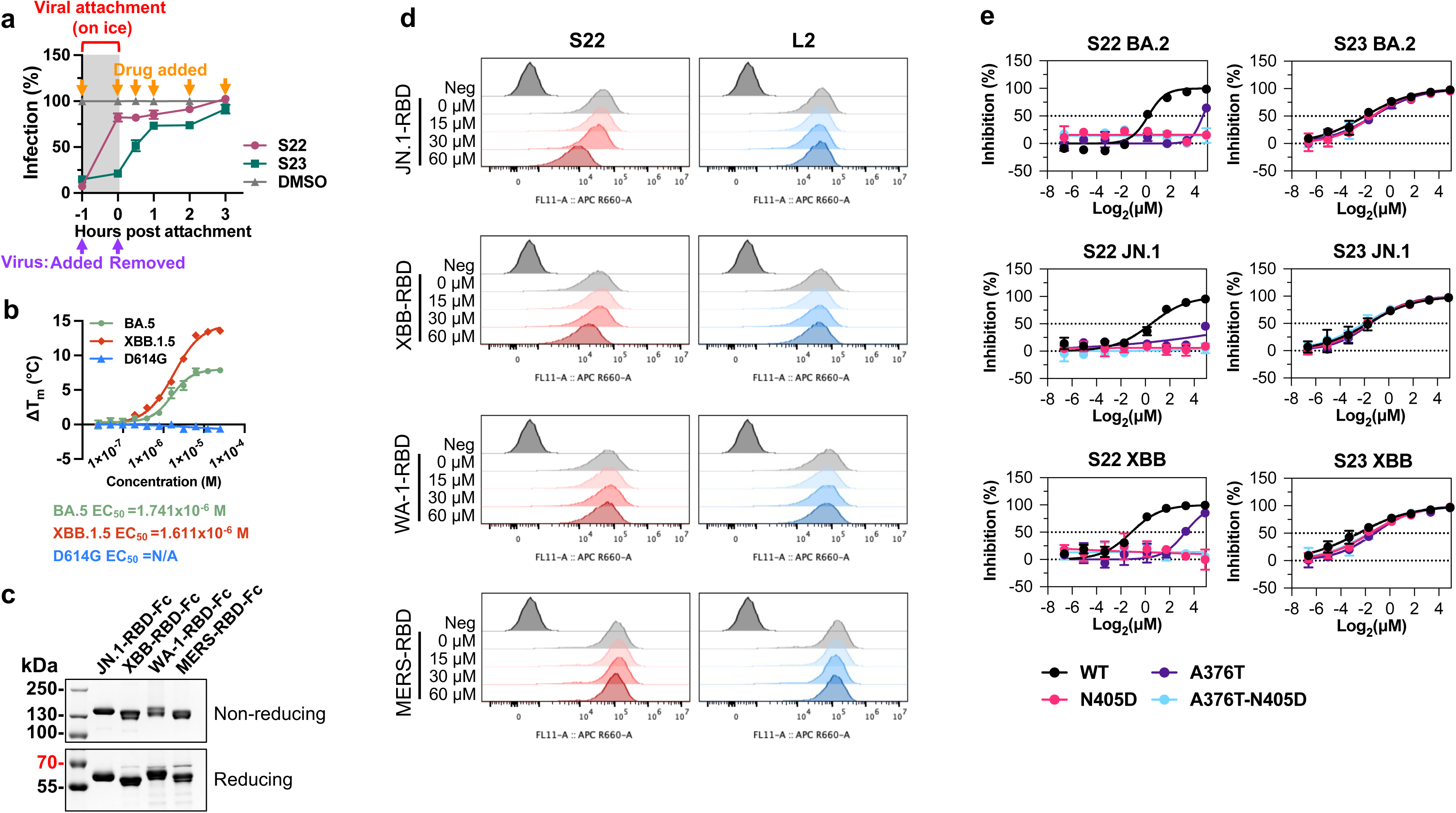
S22 acts at viral attachment by interfering with RBD-ACE2 interactions. **a**, To determine the stage of action of S22, H1299-hACE2 cells were preincubated on ice with BA.5-PVs (MOI = 2) for 1 h to allow viral attachment (from −1 h to 0 h, gray-shaded period) before medium change and transferring to 37 °C. S22 or S23 was added at their respective IC_95_ values either 1 h before (−1 h), at the same time (0 h), or at 0.5, 1, 2, or 3 h after viral attachment. Luciferase activity was measured at 48 h.p.i. Infection (%) was calculated relative to time-matched DMSO controls. Data represent the mean of two independent experiments performed in triplicate, with error bars indicating s.d. **b**, Spike ectodomains from SARS2 variants XBB.1.5, BA.5, and D614G were incubated with increasing concentrations of S22 and scanned from 25°C to 95°C with a temperature gradient of 5 °C/min. Melting temperatures (T_m_) were derived from intrinsic tryptophan fluorescence at 350 nm, and ΔT_m_ values were calculated relative to DMSO controls. Data represent mean ± s.d. EC_50_ values were determined by nonlinear regression using a variable slope four-parameter model. **c**, One microgram of purified RBD-Fc proteins was analyzed on Novex 4–20% Tris-Glycine gradient gels under non-reducing (up) and reducing (down) conditions and stained with Bio-Safe Coomassie G-250 Stain. Position and sizes of the marker proteins are indicated. **d**, The representative histograms of the results in Fig. 2a. **e**, Neutralization assays similar to Fig. 2d were conducted to evaluate S22’s activity against PVs bearing WT BA.2, JN.1, or XBB.1.5 spikes and their corresponding variants with the indicated mutations. S23 was included as a control. Data shown are from three independent experiments (n = 2 replicates per concentration), with error bars indicating s.d.

**Extended Data Fig. 4.**
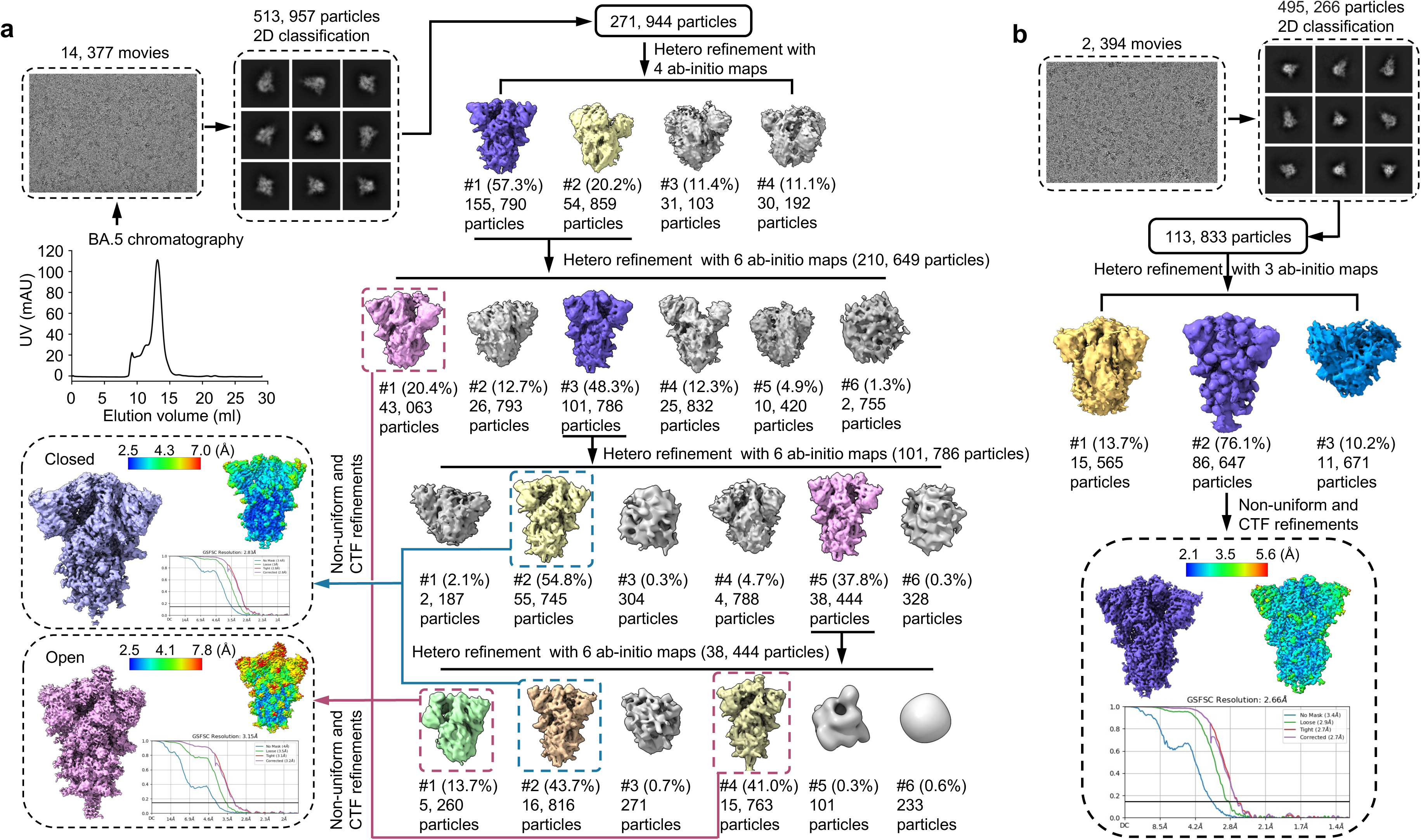
Cryo-EM analysis of the SARS-CoV-2 BA.5 spike apo structure and the BA.5 spike-S22 complex. **a**, Workflow for cryo-EM analysis of the BA.5 spike in the apo form. Size-exclusion chromatography profiles, representative cryo-EM micrographs, and 2D class averages are shown. 3D classification yielded particles with either closed or one-RBD-up conformations, which were refined to 2.83 Å and 3.15 Å, respectively. Final maps, Fourier shell correlation (FSC) curves based on the gold-standard 0.143 criterion, and local resolution estimates are shown in the black dashed boxes. **b**, Workflow for cryo-EM analysis of the BA.5 spike-S22 complex. Representative micrographs and 2D class averages are shown. 3D refinement of particles resulted in a 2.66 Å reconstruction with all RBDs in the down conformation. The final map, the associated FSC curve, and the local resolution distribution are presented in the black-dashed box.

**Extended Data Fig. 5.**
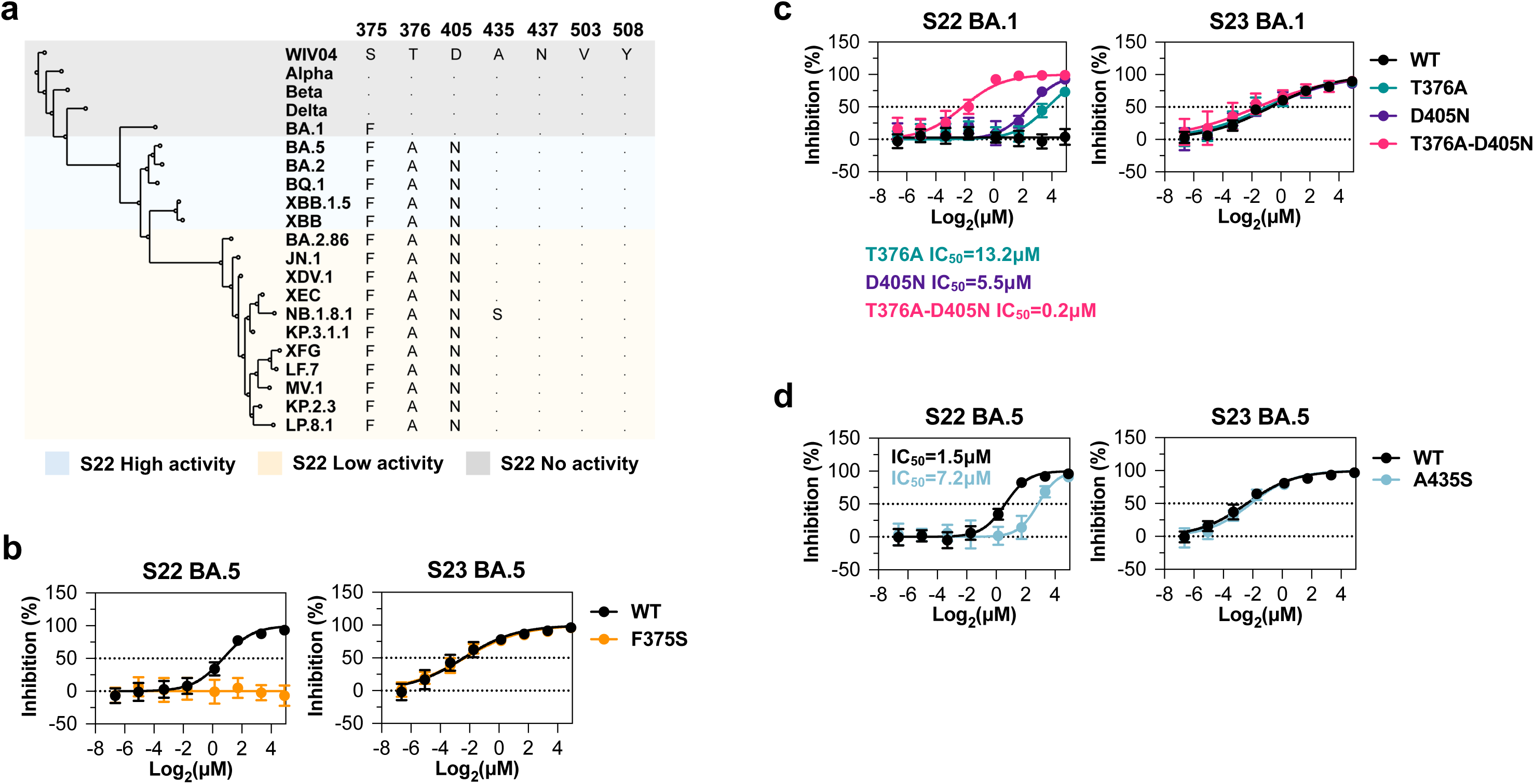
S22-interacting residues are conserved among Omicron spike variants. **a**, Phylogenetic analysis of representative SARS2 spike proteins, using WIV04 spike as the reference sequence. Alignment of residues critical for S22’s activity is shown, and the residues identical to WIV04 are indicated by dots. Shading indicates the relative sensitivity of each variant to S22, as shown in Fig. 1e. **b-d,** Neutralization assays similar to Fig. 2d were conducted to compare S22’s activity against PVs bearing WT BA.5 and a BA.5 mutant with the F375S mutation (**b**), WT BA.1 and BA.1 mutants with the indicated substitutions (**c**), and WT BA.5 and BA.5 containing the A435S mutation (**d**). S23 was included as a control. Data shown are from three independent experiments (n = 2 replicates per concentration), with error bars indicating s.d. Selected IC_50_ values are reported in the figures and are color-coded for comparison.

**Extended Data Fig. 6.**
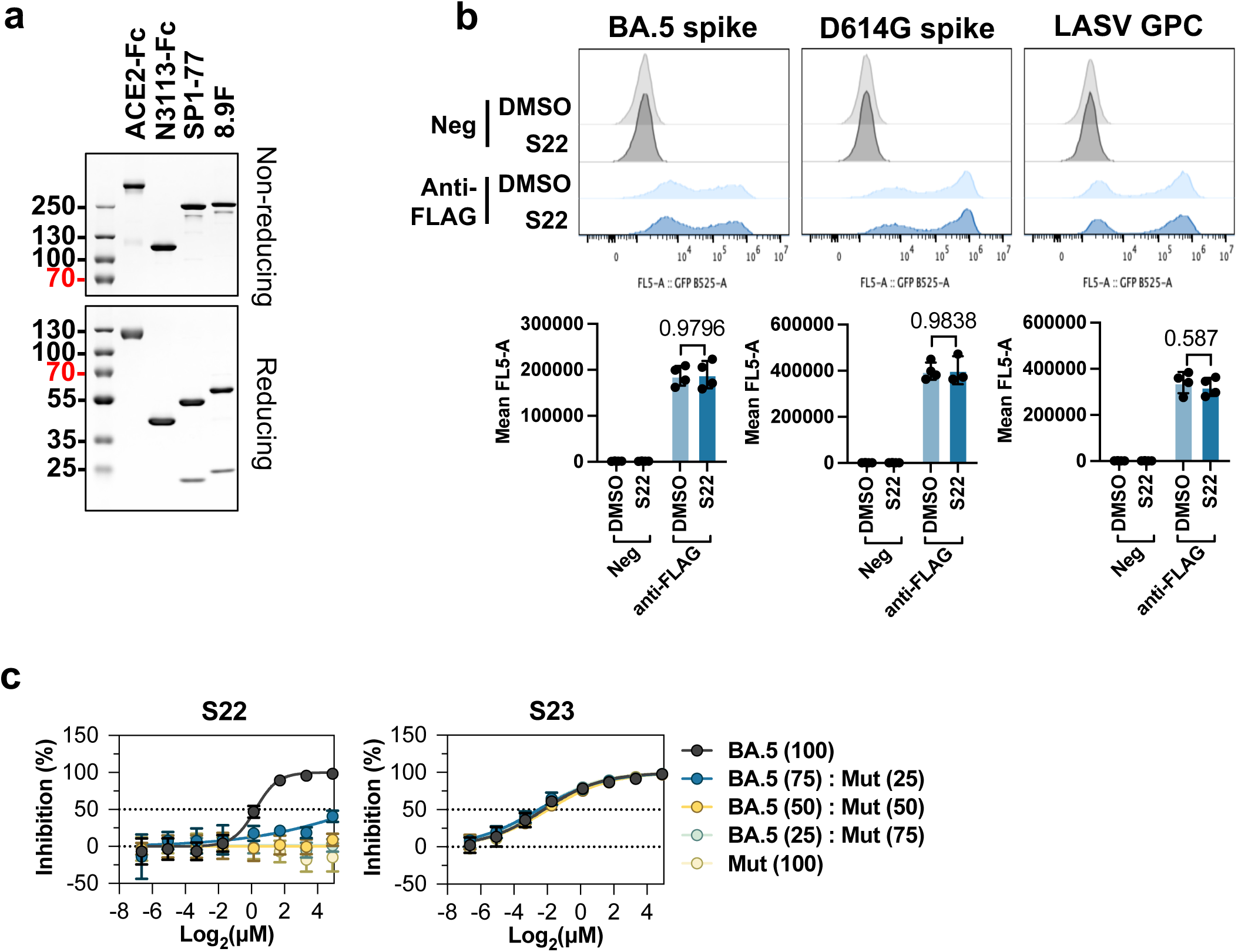
S22’s neutralizing activity requires engagement of all three RBDs in the spike trimer. **a**, Purified ACE2-Fc, N3113-Fc, SP1-77, and 8.9F used in Fig. 4 were analyzed by SDS-PAGE. **b**, Expression of the indicated entry proteins in HEK293T cells after S22 or DMSO treatment, shown in Fig. 4a, was measured by staining the C-terminal FLAG tag in the permeabilized cells and analyzed using flow cytometry. Bars show MFI from four independent experiments, and error bars indicate s.d. MFI values were square-root-transformed and analyzed using two-way ANOVA, followed by Sidak’s multiple comparisons test. Adjusted *p*-values are reported. **c**, S22’s activity was assessed against PVs produced from cells transfected with WT BA.5 spike (BA.5), the S22-resistant BA.5 F375S mutant (Mut), or their mixtures at the indicated ratios shown in brackets. S23 was included as a control. Data shown are from three independent experiments (n = 2 replicates per concentration), with error bars indicating s.d.

**Extended Data Fig. 7.**
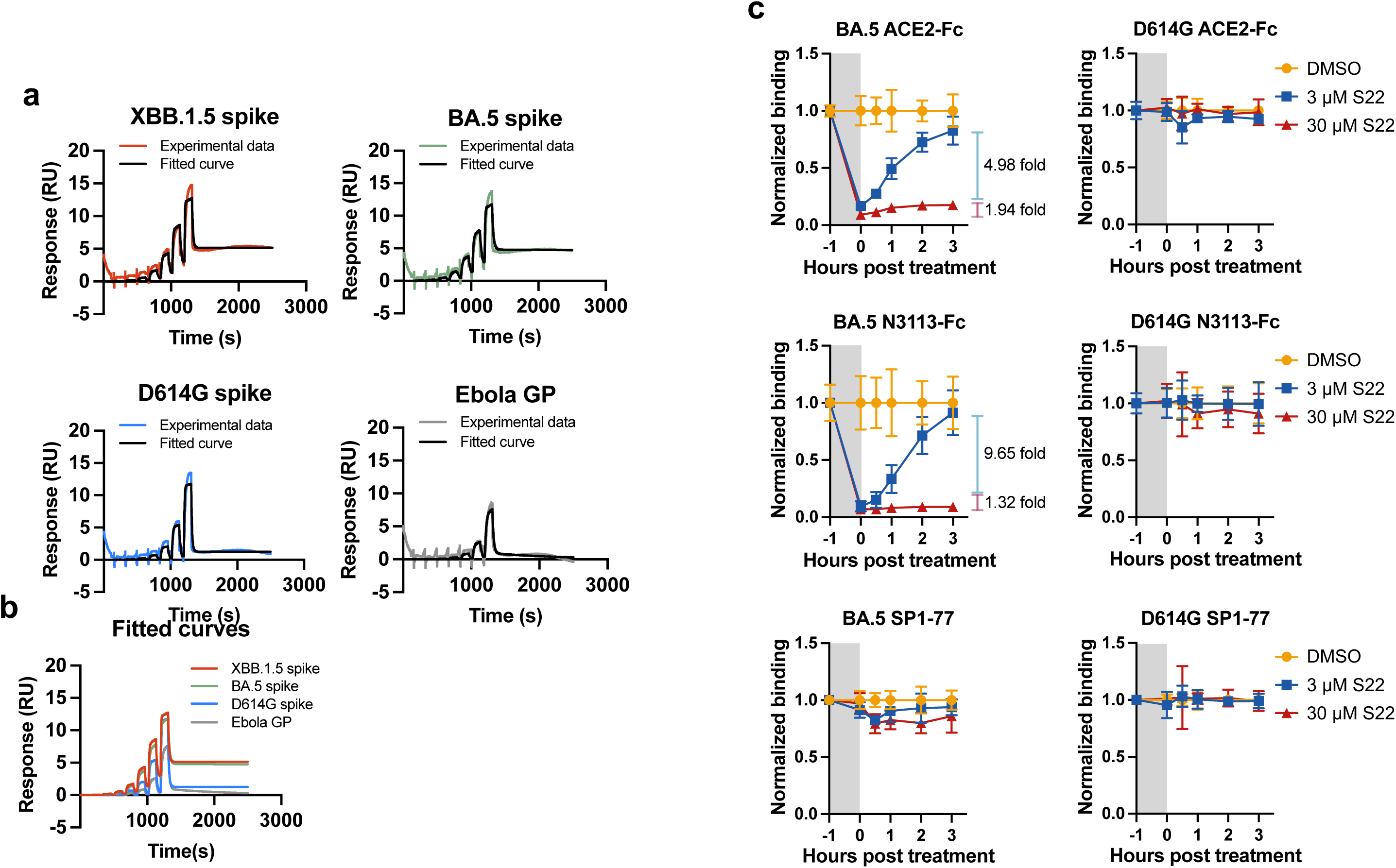
Trimeric S22 dissociates slowly from Omicron spike proteins once bound. **a**, To assess S22 binding to spike proteins, a threefold serial dilution of S22 (7.6 nM to 50 μM) was injected over immobilized SARS2 spike ectodomains (XBB.1.5, BA.5, and D614G) or EBOV GP (control) at 25 °C. Sensorgrams were reference-subtracted and globally fitted using a 1:1 Langmuir model. Experimental data are shown as colored curves, and global fits are shown as black curves. Data are representative of two independent experiments. **b**, Overlay of fitted sensorgrams from **a** shows notably different baseline signal levels between S22-binding and non-binding spikes. **c**, To assess the binding kinetics of S22 to the BA.5 spike, HEK293T cells expressing the BA.5 spike, including D614G spike as a control, were treated with S22 (3 or 30 μM) or DMSO at 37 °C for 1 h (−1 to 0 h, grey shaded area), followed by medium replacement. Cells were fixed before treatment (−1 h), at the completion of treatment (0 h), and at 0.5, 1, 2, and 3 h post-wash, then stained with ACE2-Fc and the indicated antibodies, followed by APC-conjugated anti-human Fc antibody. Data shown are representative of three independent experiments with error bars indicating s.d.

**Extended Data Fig. 8.**
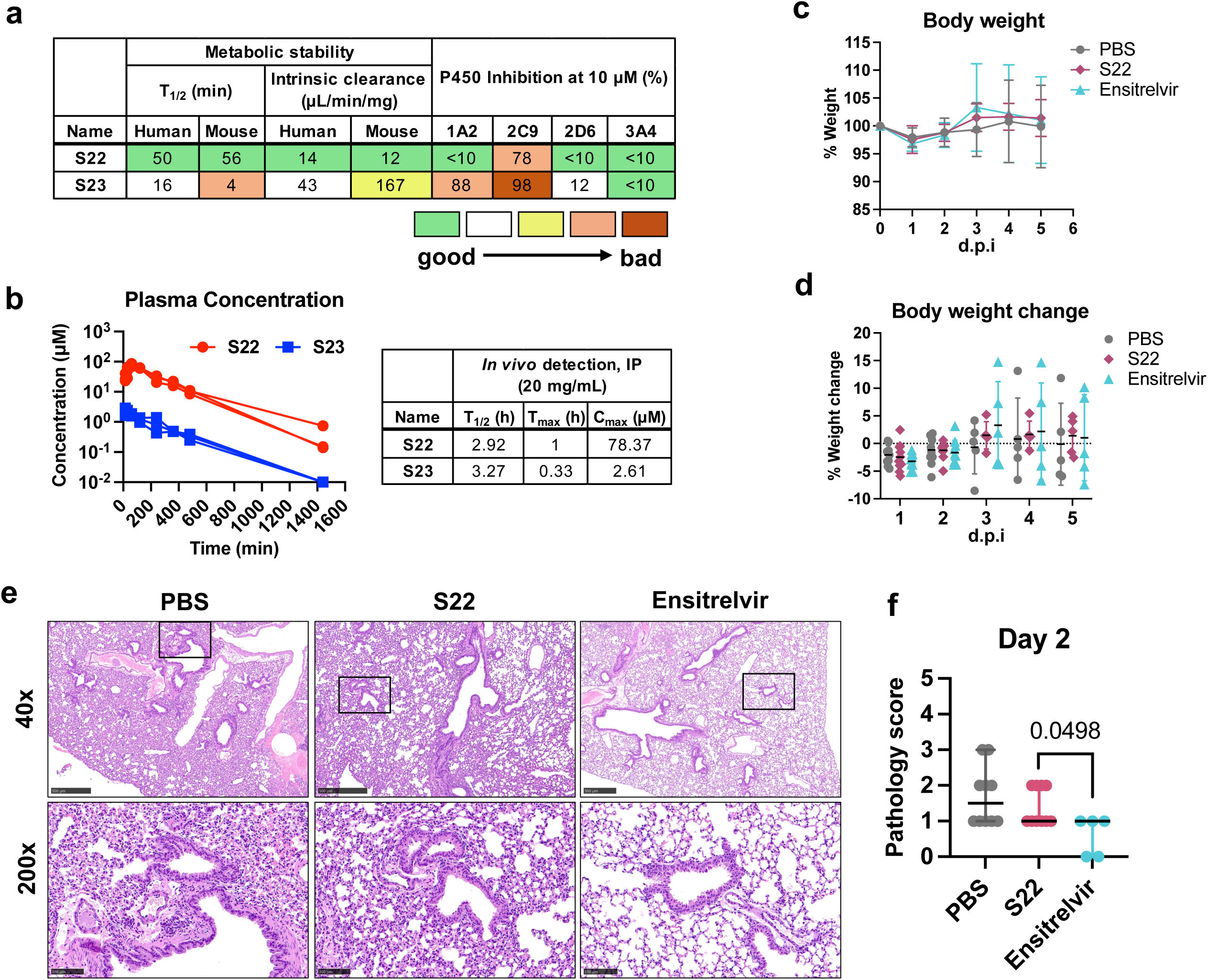
Metabolic stability, pharmacokinetics, and in vivo antiviral efficacy of S22. **a**, In vitro DMPK properties of S22 and S23, color-coded from green (good) to brown (bad). **b**, Single-dose pharmacokinetics of S22 and S23 in female C57BL/6J mice (n=3) following intraperitoneal administration at 20 mg/kg. Plasma concentrations were measured at the indicated time points by LC-MS/MS. Pharmacokinetic parameters, including maximum plasma concentration (C_max_, µM), time to reach maximum concentration (T_max_, h), and elimination half-life (T_1/2_, h), are summarized in the accompanying table. **c,d**, Body weight (**c**) and percent weight change (**d**) of mice following SARS2 infection and treatment with S22, ensitrelvir, or PBS. Data are presented as mean ± s.d. **e,f**, Representative H&E-stained lung sections at both 40x and 200x magnifications (**e**) and corresponding pathology scores (**f**) at two d.p.i. Day 5 images and pathology scores are shown in Fig. 5c**,d**. Scale bars: 500 µm for 40× images; 100 µm for 200× images. Pathology scores were evaluated by ordinal logistic regression model, with significance determined by chi-square tests with Pearson’s correction. Adjusted *p*-values are reported only for comparisons reaching statistical significance.

## Methods

### Ethics statement

Mouse studies were performed in accordance with institutional and national guidelines and were approved by the Institutional Animal Care and Use Committees (IACUC) at UF Scripps Biomedical Research (protocol 15-022) and at the University of Louisville (protocol 22134). Both in vivo pharmacokinetics and antiviral efficacy studies were evaluated in 8- to 10-week-old female C57BL/6 mice (Jackson Laboratories, strain 000664). Mice housed at UF Scripps were used for pharmacokinetic evaluations of candidate compounds, and those housed at the University of Louisville were used for antiviral efficacy studies.

### Compounds

Information for the 21 SARS-CoV-2 (SARS2) candidate compounds, as well as the control compounds, L2, M8, hydroxychloroquine (HCQ), and ensitrelvir, including sample ID, chemical structure, SMILES, PubChem CID, vendor and catalog number, molecular weight, antiviral potency, and cytotoxicity, is summarized in **Extended Data Table 1**. For in vitro experiments, all compounds were dissolved in 100% dimethyl sulfoxide (DMSO; Sigma-Aldrich, D2650) to prepare 10 mM stock solutions. For in vivo studies, S22 and ensitrelvir were formulated in 5% (v/v) DMSO, 5% (v/v) Tween-80 (Sigma-Aldrich, 655207), and 90% (v/v) sterile water.

### Cells

Expi293F cells (Gibco, A14527) were cultured in Expi293F Expression Medium (Gibco, A1435101) at 37°C in a shaking incubator at 125 rpm and 8% CO_2_. HEK293T cells (ATCC, CRL-3216) were maintained in Dulbecco’s Modified Eagle Medium (DMEM; Gibco, 10569010) supplemented with 10% (v/v) Fetal Bovine Serum (FBS; Gibco, A5670701), 1% (v/v) MEM Non-Essential Amino Acids (NEAA; Gibco, 11140-050). Vero E6 cells (ATCC, CRL-1586) were grown in Advanced MEM (Gibco, 12492013) supplemented with 10% FBS and GlutaMAX Supplement (Gibco, 35050061). Vero E6-hACE2 cells were grown in DMEM containing 10% FBS and 2 mM L-glutamine (Gibco, 25030081)^46^. H1299-hACE2 cells were maintained in RPMI 1640 Medium (Gibco, 61870036) supplemented with 10% FBS, 1% (v/v) sodium pyruvate (Gibco, 11360070), and 1 μg/mL of puromycin (Sigma-Aldrich, P9620), and A549-hACE2 cells were cultured in DMEM/F-12 (Gibco, 11320033) supplemented with 10% FBS, 1% NEAA, and 1 μg/mL of puromycin^23,47^. H1299-hACE2 cells stably expressing human dipeptidyl peptidase 4 (H1299-hACE2-hDPP4) were produced by transducing H1299-hACE2 cells with vesicular stomatitis virus (VSV) G protein-pseudotyped murine leukemia viruses (MLV) containing pQCXIB-hDPP4-FLAG, as described previously^48^. Transduced cells were selected and maintained with RPMI 1640 Medium supplemented with 10% FBS, 1% sodium pyruvate, 1 μg/mL of puromycin, and 10 μg/mL blasticidin (InvivoGen, ant-bl-1). hDPP4 expression in these cells was confirmed by cell staining using a monoclonal anti-FLAG M2-FITC antibody (Sigma-Aldrich, F4049). Except for Expi293F cells, all other cells were maintained in medium also supplemented with 1% penicillin/streptomycin (Gibco, 15140122) and cultured at 37°C in 5% CO_2_ unless otherwise specified.

### Live SARS-CoV-2 viruses

SARS2 ancestral strain WA-1 (nCoV/Washington/1/2020) and Omicron strains BA.5 (hCoV-19/USA/COR-22-063113/2022), XBB.1.5 (hCoV-19/USA/MD-HP40900/2022), JN.1 (hCoV-19/USA/New York/PV96109/2023), KP.2 (hCoV-19/USA/CA-GBW-GKISBBBB26982/2024), and KP.3 (hCoV-19/USA/NJ-GBW-GKISBBBB88291/2024) were obtained from BEI Resources, National Institute of Allergy and Infectious Diseases (NIAID).

### Protein production and purification

The immunoadhesin forms of coronavirus receptor-binding domains (RBD-Fcs) and soluble human ACE2 were produced as described previously^48^. The SARS2 nanobody N3113^49^ expression plasmid was generated by ligating synthesized gBlocks (IDT) in frame with the human IgG1 Fc domain into the pCAGGS vector using the In-fusion Snap Assembly system (Takara, 638948). The resulting constructs were transiently transfected into Expi293F cells using PEI MAX (Kyfora Bio, 24765). For antibody production, the heavy (H) and light (L) chains of the SARS2 antibody SP1-77 and the LASV antibody 8.9F were cloned into the pCMVR-VRC01-H and -L plasmids, respectively, as described previously^50^. Expi293F cells were co-transfected with H- and L-chain plasmids at a 2:1 ratio. Four to five days post-transfection (d.p.t.), culture supernatants were collected, clarified by centrifugation, and filtered through 0.22 μm filters. Fc-fusion proteins and antibodies were purified using HiTrap MabSelect Sure cartridges (Cytiva, 11-0034-93) following the manufacturer’s instructions, and eluates were buffer-exchanged into PBS and concentrated using Amicon Ultra Centrifugal Filters (Millipore, UFC9030). Purified proteins were assessed by electrophoresis on 4–20% Novex™ Tris-Glycine gels (Invitrogen) under both reducing and non-reducing conditions and stained by Coomassie blue (Bio-Rad, 16107787).

Plasmids encoding the ectodomains of SARS2 D614G spike, BA.5 spike, XBB.1.5 spike, and Ebola virus glycoprotein were constructed by subcloning the corresponding coding sequences into the pCAGGS vector with a C-terminal T4 foldon trimerization domain and His tag for purification. To stabilize the SARS2 spike ectodomain in the prefusion conformation, several mutations were introduced, including two mutations at the furin cleavage site (RRAR to AGAR) and six proline substitutions within the S2 subunit^51^. To produce these proteins, Expi293F cells were transiently transfected with the corresponding plasmids at a density of 3 × 10^6^ cells/mL using PEI MAX. Three d.p.t., supernatants were harvested and buffer-exchanged into PBS. The spike or GP proteins were first purified using a Ni-NTA affinity column (Cytiva, 17524801) with elution at 500 mM imidazole and subsequently purified by size-exclusion chromatography (SEC) on a Superose 6 Increase 10/300 GL column (Cytiva, 29091596) using an ÄKTA Pure system (Cytiva). Plasmid information is shown in the **Extended Data Table 2**.

### Pseudotyped virus neutralization and cytotoxicity assays

The plasmids encoding viral entry proteins used for pseudotyped virus (PV) production are listed in **Extended Data Table 2**. PVs bearing coronavirus spikes (including mutants) and the glycoproteins of Lassa virus and Machupo virus were produced as described previously^48^. Neutralization assays were performed using the pre-titrated PVs and the cell lines H1299-hACE2, A549-hACE2, or H1299-hACE2-hDPP4. Briefly, serially diluted compounds were preincubated with diluted PVs at 37°C for 1 hour, and the compound-virus mixtures were added to cells seeded in 96-well plates (Corning, 3596). Following a 48-hour incubation at 37°C, firefly luciferase expression was determined using the Britelite plus Reporter Gene Assay System (Revvity, 6066761). Cytotoxicity was evaluated in parallel with uninfected cells under identical compound treatment conditions, using the CellTiter-Glo 2.0 Cell Viability Assay (Promega, G9242). Each experiment was performed in duplicate, and luminescence signals were measured using a VICTOR Nivo plate reader (PerkinElmer).

The percentage of viral inhibition was calculated by comparing background-subtracted luminescence signals from compound-treated infected wells with those from vehicle-treated infected control wells, which represent 100% infection. Cytotoxicity was determined by comparing background-subtracted signals from compound-treated wells with those from vehicle-treated controls. The half-maximal inhibitory (IC_50_) and cytotoxic (CC_50_) concentrations were obtained by fitting the inhibition and viability curves using nonlinear regression with a nonlinear regression method with a four-parameter logistic model in GraphPad Prism (v10.4.2).

### Live SARS-CoV-2 microneutralization assay

All in vitro work with live SARS2 was conducted within a biosafety level 2+ facility, which was approved by the Institutional Biosafety Committee at the University of Louisville. The neutralizing potency of S22 and control compounds S23 and HCQ against the live SARS2 variants was evaluated using a virus microneutralization assay, as described previously^52^. Briefly, two-fold serially diluted compounds were pre-incubated with indicated SARS2 viruses (MOI=0.01) at 37°C for 1 hour, and then the compound-virus mixtures were added to Vero E6-hACE2 cells pre-seeded in 96-well plates one day prior to infection. 96 hour post infection (h.p.i.), cell viability was determined using Neutral Red assay (Sigma-Aldrich, TOX4). The antiviral activity of each compound against the indicated SARS2 strains was determined by quantifying the concentration that reduced virus-induced cytopathic effect (CPE) by 50% (IC_50_) compared with the untreated virus control.

### Time-of-addition assay

H1299-hACE2 cells were seeded in 96-well plates one day prior to infection. On the following day, cells were pre-incubated with PVs bearing the BA.5 spike (MOI=2) on ice for 1 hour to allow viral attachment, after which the media was refreshed. The plates were then transferred to 37°C to allow viral internalization. S22 and S23 were added at their respective IC_95_s at the indicated time points relative to virus attachment: pre-attachment (added simultaneously with PV on ice, −1 h), end-attachment (added immediately after viral removal and switched to 37°C, 0 h), or post-attachment (added at 0.5, 1, 2, or 3 hours post-attachment). Luciferase activity was measured 48 h.p.i. using Britelite Plus substrate, and luminescence signals were recorded with a VICTOR Nivo plate reader. Infection (%) was calculated as the luciferase signal from each condition relative to its time-matched DMSO-treated control.

### S22 inhibition of RBD-ACE2 interaction

To assess whether S22 blocks SARS2-RBD-ACE2 interaction, purified RBD-Fc proteins (0.2 μg/mL) were preincubated with serially diluted S22, L2, or vehicle at room temperature for 30 minutes. H1299-hACE2-hDPP4 cells were detached using Accutase (STEMCELL, 07920), fixed with ice-cold 3.7% formaldehyde solution (Sigma-Aldrich, 252549), and blocked with 10% (v/v) goat serum (Gibco, 16210064) in PBS. Cells were incubated with RBD-compound mixtures on ice for 1 hour, and RBD binding was detected using goat anti-human Fc-APC antibody (Jackson ImmunoResearch, 109-135-098). All samples were analyzed on the CytoFlex S flow cytometer (Beckman Coulter), and data were processed using FlowJo v10.10 software.

### Measurement of RBD in the up or down conformation

To evaluate S22-induced conformational changes in BA.5 spike, HEK293T cells were transfected with plasmids expressing C-terminal FLAG-tagged BA.5 spike, D614G spike, or LASV GPC using PEI MAX, with empty vector as a negative control. At 48 hours post-transfection (h.p.t.), cells were treated with 30 μM S22 or DMSO vehicle at 37°C for 1 hour prior to detachment and fixation. To measure protein or antibody binding to the cell-surface spike or GPC, fixed cells were incubated with ACE2-Fc (1 μg/mL), N3113-Fc (5 μg/mL), or SP1-77 (1 μg/mL), followed by goat anti-human Fc-APC. To quantify the total expression of spike or GPC, fixed cells were permeabilized with 0.5% Triton X-100 (Sigma-Aldrich, 93443) for 10 minutes at room temperature, stained with 1 μg/mL anti-FLAG M2 antibody (Sigma-Aldrich, F1804), and detected with goat anti-mouse IgG-FITC (Jackson ImmunoResearch, 115-096-072).

To investigate the role of S22 trimerization in locking the BA.5 spike in the closed conformation, HEK293T cells were transfected with wild-type (WT) BA.5 spike, the S22-resistant BA.5 F375S mutant, or mixtures of both at the indicated ratios. At 48 h.p.t., cells were treated with S22 (3, 10, or 30 µM) or DMSO, fixed, and stained with ACE2-Fc, N3113-Fc, or SP1-77, followed by goat anti-human Fc-APC.

To explore S22-binding kinetics, HEK293T cells expressing BA.5 spike or D614G were treated with 3 μM S22, 30 μM S22, or DMSO vehicle at 37°C for 1 hour. After compound washout and medium replacement, cells were collected and fixed at the indicated time points and then stained with ACE2-Fc, N3113-Fc, or SP1-77, followed by goat-anti-human Fc-APC. All samples were analyzed on a CytoFlex S flow cytometer (Beckman Coulter), and data were processed using FlowJo v10.10 software.

### Cryo-EM structural study

For cryo-EM analysis, BA.5 spike ectodomain (0.8 mg/mL, see the Protein production and purification section), either alone or mixed with 100 μM S22, was applied to freshly glow-discharged Quantifoil R1.2/1.3 300-mesh copper grids (Electron Microscopy Sciences). 4 µL of each sample was deposited onto grids, blotted for 3.5 s at 22°C under 100% humidity, and plunge-frozen in liquid ethane using a Vitrobot Mark IV (FEI). Images were collected at the Hormel Institute, University of Minnesota, on a Titan Krios transmission electron microscope (FEI) operating at 300 kV and equipped with a BioContinuum energy filter (slit width 20 eV) and K3 direct electron detector (Gatan, Inc.) in CDS mode. Data collection was performed using the EPU software (Thermo Scientific) at a pixel size of 0.664 Å (nominal magnification 130,000x) and a defocus range of −2.5 to −0.75 μm. A total of 14377 movies were collected for the apo BA.5 spike (total electron dose 52 e^−^/Å^2^), and 2394 movies for BA.5 spike-S22 complex (total electron dose 50 e^−^/Å^2^). Cryo-EM data collection parameters are summarized in **Extended Data Table 3**.

### Cryo-EM data processing, model building, and refinement

Cryo-EM data were processed using cryoSPARC v4.5.1^53^, following the workflow shown in **Extended Data Fig. 4**. Beam-induced motion was corrected using MotionCor2^54^, and contrast transfer function (CTF) parameters were estimated with CTFFIND-4.1.13^55^. Micrographs with CTF fits worse than 7 Å were excluded. Particle picking was performed using the Blob picker followed by Template-based picking in cryoSPARC v4.5.1. Three rounds of 2D classification were used to remove junk particles, and particles from well-defined 2D classes were subjected to *ab initio* reconstruction and heterogeneous refinement. For the apo BA.5 spike dataset, four rounds of 3D classification revealed two conformations: a closed spike with all three RBDs down and an open spike with one RBD up. Particles corresponding to each conformation were refined separately using non-uniform refinement and CTF refinement to obtain the final maps. For the BA.5 spike-S22 complex, a single predominant conformation with all RBDs down was observed after one round of 3D classification, and particles from selected classes were refined using the same refinement strategy. Final map resolution was estimated using the gold-standard Fourier shell correlation (FSC) at 0.143 between half-maps, and local resolution was calculated in cryoSPARC v4.5.1. Initial model building of the BA.5 spike-S22 complex was performed in Coot-0.8.9^56^ using the BA.4 spike structure (PDB: 8CIN) as the starting model. Additional densities were observed in the central cavity of the spike trimer and were fitted with three S22 molecules. Iterative model refinement was performed using Phenix-1.16^57^ and manual adjustment in Coot-0.8.9. Model statistics are summarized in **Extended Data Table 3**. Structural figures were generated using UCSF Chimera X v0.93^58^ and PyMOL v3.0 (Schrödinger, LLC), and contact residues at the BA.5 spike-S22 interface were analyzed using LigPlot^+^ V.2.3.

### Surface plasmon resonance

Single-cycle kinetics analyses of S22 binding to SARS2 spike proteins or to the control protein EBOV GP were performed on a Biacore 1S+ instrument (Cytiva) at 25°C. XBB.1.5, BA.5, D614G spike or EBOV GP ectodomains (see the Protein production and purification section) were immobilized on flow cells of CM5 sensor chips (Cytiva, 29149603) to ∼5000 response units (RU) using the amine coupling method (Cytiva, BR100050), following the manufacturer’s protocol. Reference flow cells were activated and deactivated without protein coupling to serve as background controls. A threefold serial dilution of S22, ranging from 7.6 nM to 50 µM (nine concentrations in total), was injected over both the reference and sample flow cells at a flow rate of 30 µL/min in 1x HBS-EP+ buffer (Cytiva, BR100669) supplemented with 1% DMSO. Each injection included a 100 s association phase followed by a 120 s dissociation phase, which was conducted in single-cycle mode without surface regeneration. DMSO calibration was performed at the beginning and end of each run to correct for bulk refractive index changes caused by varying DMSO concentrations in the compound solutions. Reference-subtracted sensorgrams were globally fitted using Biacore Insight Evaluation Software v6.0.7.1750, using a 1:1 Langmuir binding model to derive kinetic parameters. Each experiment was independently repeated at least twice on separate immobilized surfaces, and results are reported as fitted values with 95% confidence intervals.

### Nano differential scanning fluorometry (NanoDSF)

NanoDSF was performed to evaluate S22 binding-induced thermal stabilization of SARS2 spike proteins using a Prometheus Panta instrument (NanoTemper Technologies). Purified XBB.1.5, BA.5, D614G spike and EBOV GP proteins (see the Protein production and purification section) were buffer-exchanged into 1x HBS-EP+ buffer (Cytiva) using Zeba^TM^ spin desalting columns (Thermo Scientific, 89878). A twofold serial dilution of S22 (11 concentrations ranging from 50 µM to 48 nM) was prepared in assay buffer (10 mM HEPES, 150 mM NaCl, pH 7.4) with DMSO supplemented to a final concentration of 1% and mixed at a 1:1 (v/v) ratio with spike proteins (final concentration at 0.1 mg/mL). 10 µL of each mixture was loaded into Prometheus High Sensitivity Capillaries (NanoTemper, PR-C006) and scanned from 25°C to 95°C with a temperature gradient of 5 °C/min. Intrinsic protein fluorescence at 330 nm and 350 nm was recorded. Melting temperatures (T_m_) were determined from the 350 nm signal by fitting to a two-state Boltzmann model with exponential baselines:

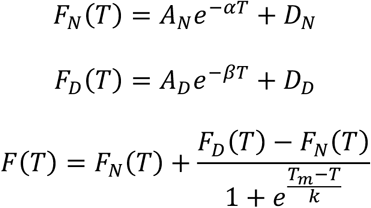

*F_N_*(*T*): Fluorescence of the native (folded) state at temperature T
*F_D_*(*T*): Fluorescence of the denatured (unfolded) state at temperature T
*F*(*T*): Observed fluorescence of both native and denatured populations at temperature T
*A_N,_ a, D_N_*: Amplitude, decay rate, and offset for the native-state baseline, respectively
*A_D,_ b, D_D_*: Amplitude, decay rate, and offset for the denatured baseline, respectively
*T_m_*: Melting temperature at which 50% of protein is unfolded
*k*: Slope factor; reflects the steepness of protein unfolding transition

Ligand-induced stabilization (ΔTₘ) was plotted against S22 concentration and fitted in GraphPad Prism using a four-parameter logistic (4PL) model to obtain EC_50_ values with 95% confidence intervals.

### Hepatic microsomal stability and cytochrome P450 inhibition assays

Microsomal stability of S22 and S23 was assessed using INVITROCYP 150-donor pooled human liver microsomes (BioIVT, X008070) and mouse liver microsomes (BioIVT, M000151). 1 µM of each test compound was incubated with 0.5 mg/mL hepatic microsomes in 100 mM potassium phosphate (KPi) buffer at pH 7.4 at 37°C. Reactions were initiated by adding NADPH (Sigma-Aldrich, N7505) to a final concentration of 1 mM. Aliquots were collected at 0, 5, 10, 20, 40, and 60 minutes, and quenched with a fivefold volume of acetonitrile (Sigma-Aldrich, 34851) to stop the reaction and precipitate microsomal proteins. Parallel incubations lacking NADPH were included to assess P450-independent compound depletion. Samples were centrifuged through Multiscreen Deep 96-Well Solvinert filter plates (Millipore, MDRPNP410), and supernatants were quantified by LC-MS/MS (SCIEX QTRAP 6500) for the remaining compounds. Data were log-transformed and represented as half-life (T_1/2_, min) and Intrinsic clearance (µL/min/mg).

To assess potential drug-drug interactions, S22 and S23 were screened for inhibition of four major human cytochrome P450 (CYP) isoforms (CYP1A2, CYP2D6, CYP2C9, and CYP3A4), using pooled human liver microsomes (BioIVT, X008070). The activities of these CYP isoforms were measured by monitoring the metabolism of selective probe substrates: phenacetin O-deethylation to acetaminophen (CYP1A2), tolbutamide hydroxylation to hydroxytolbutamide (CYP2C9), bufuralol hydroxylation to 4′-hydroxybufuralol (CYP2D6), and midazolam hydroxylation to 1′-hydroxymidazolam (CYP3A4). All the substrates and reference inhibitors used were purchased from Sigma-Aldrich. Incubations were performed at 37°C in the presence or absence of 10 µM test compound, with each substrate used at a concentration near its respective Kₘ. Specific reference inhibitors for each isoform (furafylline for CYP1A2, sulfaphenazole for CYP2C9, quinidine for CYP2D6, and ketoconazole for CYP3A4) were included in each assay to validate enzyme selectivity and system performance. Reactions were terminated by adding the ice-cold acetonitrile (3x volume, v/v) to precipitate proteins, followed by centrifugation at 3,000x g for 10 minutes. Supernatants were analyzed by LC-MS/MS to quantify metabolite formation. Percent inhibition was calculated by comparing metabolite formation rates in compound-treated samples relative to those in vehicle-treated controls.

### In vivo pharmacokinetics study in mice

In vivo pharmacokinetics of S22 and S23 were evaluated in 8- to 10-week-old female C57BL/6 mice by intraperitoneal (IP) administration of the test compound at 20 mg/kg. Compounds were formulated in a vehicle comprising 5% (v/v) DMSO, 5% (v/v) Tween-80, and 90% (v/v) sterile water. Three mice were used per compound. 25 µL of blood samples were collected into heparin-coated capillary hematocrit tubes (Drummond Scientific Company, 8-000-7520-H/5) at 0, 15, 30, 60, 120, 240, 360, 480, and 1440 minutes post-dose using a microsampling technique. Plasma was separated by centrifugation, and compound concentrations were quantified using LC-MS/MS. Pharmacokinetic parameters, including maximum plasma concentration (C_max_, µM), time to maximum concentration (T_max_, h), and elimination half-life (T_1/2_, h), were determined by non-compartmental analysis using Phoenix WinNonlin software (Pharsight Inc.).

### Inhibition of SARS-CoV-2 infection in mice

Female C57BL/6J mice (n = 10 per group, 8- to 10-week-old) were anesthetized and intranasally infected with Omicron XBB.1.5 variant in 50 µL DMEM (10^4^ PFU/mouse, 25 µL per nostril). At 4 h.p.i., mice received an intraperitoneal administration of S22 (40 mg/kg body weight), ensitrelvir (20 mg/kg body weight), or PBS in 200 µL. Body weight and clinical activity were monitored daily. Mice were euthanized on two and five days post-infection (d.p.i.), and lung tissues were collected. The left lung lobe was fixed in 10% (v/v) neutral-buffered formalin for histopathology, and the remaining tissue was homogenized in sterile PBS using a handheld tissue homogenizer (Omni International) and stored at −80°C for virological analysis.

Viral titers in lung homogenates were determined by the 50% tissue culture infectious dose (TCID_50_) assay^52^. Briefly, Vero E6 cells were seeded at 2 × 10^4^ cells per well in 96-well flat-bottomed tissue culture plates and incubated overnight before infection. Lung homogenates were clarified by centrifugation at 8,000 rpm for 10 minutes at 4°C, 10-fold serially diluted (up to 10^−7^) in infection medium (DMEM supplemented with 5% FBS and 1% antibiotic-antimycotic), and added to pre-seeded cells. Cells were fixed at 4 d.p.i. and stained with 0.1% crystal violet (Sigma-Aldrich, C0775) in 10% neutral-buffered formalin, and cytopathic effect was scored. Each experiment was performed in quadruplicate, and the mean TCID_50_ values were calculated using the Reed-Muench method^59^ and normalized to tissue weight (per gram of lung).

Fixed lung tissues were paraffin-embedded, sectioned, and stained with hematoxylin and eosin (H&E) by iHisto (Everett, MA). The HE-stained slides were examined by a board-certified veterinary pathologist using virtual microscopy software on high-resolution digital images^60,61^. Tissues were evaluated for SARS2-associated disease severity by scoring pulmonary edema (0 = none, 1 = <25%, 2 = <50%, 3 = <75%, and 4 = ≥75% of examined fields). Perivascular and peribronchiolar infiltrating lymphoid cell aggregates were scored according to a modified method adapted from previous reports^62^: 0 – within normal parameters; 1 – small, scattered cell infiltrates; 2 – Small aggregates ∼<3 cells thick; 3 – multifocal aggregates ∼>3 cells thick; 4 – large aggregates readily visible at low magnification that can compress adjacent lung tissues. The cumulative scores from these histopathological parameters were combined to generate a “pathology score” for each mouse lung.

To monitor spike escape mutations in virus populations in drug-treated mice, lung homogenates and XBB.1.5 virus were treated with TRIzol reagent (Invitrogen, 15596026), and total RNA was extracted using the Direct-zol RNA Miniprep Plus Kit (Zymo Research, R2070). First-strand cDNA was synthesized from 1 µg of total RNA using the SuperScript IV First-Strand Synthesis System (Invitrogen, 18091200), according to the manufacturer’s instructions. The spike gene was amplified from cDNA by nested PCR using PrimeSTAR Max DNA Polymerase (Takara, R045A), with the second-round primers containing sample-specific barcodes and unique molecular identifiers (UMIs). PCR products were gel-purified using the NucleoSpin Gel and PCR Clean-up Kit (Takara, 740611), and subjected to Oxford Nanopore Sequencing (Plasmidsaurus). Second-round primer sequences used are listed in **Extended Data Table 4**.

### Bioinformatics

The drug-like properties of S22 and S23 were predicted using the SwissADME server (www.swissadme.ch). The alignment of the 10 SARS2 spike sequences shown in **Fig. 2b** was performed using Geneious Prime 2025 (Dotmatics). Phylogenetic analysis of SARS2 spikes was conducted by aligning spike protein sequences from representative SARS2 variants using MUSCLE3^63^ under default parameters. A phylogenetic tree was generated from the aligned sequences using the neighbor-joining method, and rerooted to the WIV04 reference strain. The final tree was visualized with the ete3 Python package. The Python script used for sequence alignment, tree construction, and generation of the mutation summary table (**Extended Data Fig. 5a**) is included as “SARS2 Phylogeny Code.zip” in the Supplementary Information. Multiplexed nanopore sequencing of the spike protein ORF was performed as described above (see the “Inhibition of SARS2 infection in mice” section), using hexanucleotide-barcoded forward primers to differentiate samples. Sequences were processed using an in-house C++ program, included as “Spike Escape Mutation Analysis.zip” in the Supplementary Information, that parsed *.fastq* output, filtered invalid reads, and aligned the outputs at the amino acid level to the reference XBB.1.5 spike (EPI_ISL_16292655). Specifically, reads were excluded from analysis if they (1) contained any N characters, (2) did not contain the oligonucleotide sequences CTTGTTAACAACTAAACGAACAATG and ACGAACTTATGGATTTGTTT which flank the ORF at the 5’ and 3’, respectively, (3) lacked recognizable barcodes, (4) were less than 2610 nucleotides long, (5) contained premature stop codons, or (6) did not end with a stop codon. Qualified reads were aligned to the reference using the Needleman-Wunsch algorithm and the BLOSUM62 scoring matrix. These alignments were used to tally the frequencies of amino acid substitutions and deletions relative to the reference sequence as reported in **Fig. 5e**.

### Statistics

Data normality and homoscedasticity were assessed using residual plots (GraphPad Prism version 10.0.0 for Windows, GraphPad Software, Boston, Massachusetts USA, www.graphpad.com”). Binding assay mean values were square-root transformed prior to two-way ANOVA to achieve normally distributed residuals. IC50 values were analyzed on the natural log scale for ANOVA. Lung viral load values were log10-transformed prior to mixed model ANOVA. When log10 or square root transformations failed to satisfy these assumptions, we employed generalized linear model with gamma distribution and log link function (2025 JMP® 18. JMP Statistical Discovery LLC, Cary, NC). All tests were two-sided with α = 0.05.

Factorial models and post hoc comparisons: For one- and two-way designs, fixed effects such as treatment, strain/mutation, dose, spike mix, day post infection, and their interactions as appropriate for each experiment. Interaction terms were retained when statistically significant; otherwise, main-effects models were used. Post hoc comparisons were conducted only when treatment effects were statistically significant, except for pre-planned comparisons. Multiple comparison corrections were applied as appropriate: Tukey’s HSD was used for comparisons among all groups, Dunnett’s test was used for control comparisons within a treatment group, and Sidak’s test was used for planned pairwise contrasts. GLM model was tested on JMP and Tukey-HSD was used for multiple testing correction.

Dose-response modeling for neutralization assays: Nonlinear regression method was used to fit curves with 4-parameter logistic model to the dose-response datasets in GraphPad Prism. If the fit failed due to non-identifiable parameters, unstable Hill slope, if less than 50% inhibition was observed across the tested concentration range or if the Hill slope was negative, the drug/protein dataset was categorized as “ no detectable inhibition”. When a successful fit was achieved, natural log-transformed IC50 values were compared using 95% CI if there was one experiment with multiple replications, lognormal ANOVA was used when the experiment was repeated 3 times with 4 replications per group. For neutralization assays, samples achieving ≥50% inhibition at any tested concentration were classified as “responders” and those failing to reach 50% inhibition were classified as “non-responders.” Differences in responder rates between groups were tested using two-sided Fisher’s exact test.

Pathology: Pathology scores (ranging from 0 to 3) were compared across two categorical variables (treatment and time), each with three and two levels, respectively. Statistical analysis was performed using ordinal logistic regression (2025 JMP® 18. JMP Statistical Discovery LLC, Cary, NC), treating pathology score as the response variable and both categorical variables as predictors. Ordinal pathology scores per day were compared across paired drugs using Chi-square tests with Pearson’s correction.

## Acknowledgements

This work was supported by the National Institute of Allergy and Infectious Diseases (NIAID) grants U19-AI171954 AND U19-AI171443. M.B.F. is the Alicia and Yaya Professor of Viral Pathogen Research at the University of Maryland School of Medicine. D.K.M. is Director of the Division of Comparative Pathology, Roy J. and Lucille A. Carver College of Medicine at the University of Iowa. R.S.H. is an Investigator of the Howard Hughes Medical Institute and the Ewing Halsell President’s Council Distinguished Chair at the University of Texas San Antonio.

## Author Contributions

H.M., B.G., M.F., and H.C. conceived the study. H.M., B.G., G.Y., D.S., J.V., J.Z., M.F., F.L., and H.C. designed experiments. R.S.H., M.D.C., T.D.B., J.Z., M.F., F.L., and H.C. supervised the study. H.M., B.G., G.Y., L.Z., D.S., F.B., J.V., S.Z., C.R.B., L.L., Y.G., J.B., C.E.K., L.B., and D.K.M. performed experiments and/or analyzed data. H.M., C.C.B., and G.C.C. performed bioinformatics analyses. G.C.C. conducted statistical analyses. H.L., Y.O., C.W., S.W., M.B.F., L.S., and T.P.S. developed key reagents and/or provided useful insights. H.M., B.G., M.F., and H.C. wrote the paper, and all authors reviewed and edited the manuscript.

